# The allometry of spiracles and gill slits: a window into the respiratory capacity of rays

**DOI:** 10.64898/2026.09.29.755487

**Authors:** Rachel M. Aitchison, Wade J. VanderWright, Amanda E. Arnold, Anthony S. Iliou, Brogan Neufeld, Logan Vuorensivu, Sophie R. Brian, Owen Kui, Jay H. Matsushiba, Olivia S. Wynn Dulvy, Sebastian I. Wynn Dulvy, Nicholas K. Dulvy

## Abstract

Metabolic rate governs the pace of life through the allocation and expenditure of energy, yet metabolic rate data are unavailable for most species. External morphological traits, such as gill slits, have been used to infer respiratory physiology and metabolism in sharks, yet, whether that pattern is generalizable across rays is unknown. In rays (Batoidea), spiracle length seems adaptive to oxygen uptake capacity, environmental oxygen availability, and activity level. Thus, we evaluate the allometry of external respiratory morphology (spiracles and gill slits) in rays using 550 museum specimens representing 88 species. We test whether spiracle length and gill slit height grow isometrically with body size (total length or disc width) within and across species. Overall, we find spiracles and gill slits are morphologically diverse, varying widely in shape within and between families. Spiracle length and gill slit height scale isometrically and are a consistent proportion of body size, despite such high morphological variability. While that proportion is consistent through ontogeny, each species proportion differs (e.g. gill slit height ranges from 0.93% of total length in Spotted Legskate to 2.93% in Pacific Electric Ray). Of the three swimming styles, pectoral oscillating species (Mobulid and Eagle Rays) may have steeper scaling of spiracle length with gill slit height–consistent with greater respiratory capacity in species with more active lifestyles. Our findings are consistent with literature hypothesizing external respiratory morphology (spiracles and gill slits) is tightly related to metabolic rate and activity level, and demonstrate the value of natural history collections in ecological research. Importantly, the isometry of both respiratory traits means that a single measure of any individual, irrespective of size, will be representative of the species thus providing a basis for expanding our understanding of metabolic morphology, enabling future work on life history, extinction risk, and ecological function.

## Introduction

Metabolic rate, or the rate at which organisms acquire, utilise, and expend energy, is considered the most fundamental biological rate (Brown et al., 2004). Metabolic scaling, the rate at which energy use increases with body mass (Kleiber, 1947), underpins large-scale ecological patterns ranging from population growth to community assembly and ecological function due to energy allocation (Brandl et al., 2023; Brown et al., 2004). Thus, metabolic rate sets the “pace of life” (Brown et al., 2004; Pettersen et al., 2016; Ricklefs & Wikelski, 2002) and is considered an ecological “super trait” (Brandl et al., 2023).

Direct measures of metabolic rate often require controlled field or laboratory settings, resulting in data for relatively few species based on a narrow, and biased, range of body sizes (Bigman et al., 2021). This is problematic for species that grow continuously throughout their lifetime (Bigman et al., 2021). Globally, there are field metabolic rate measurements for only ∼600 animal species across endotherms, ectotherms, invertebrates and vertebrates (de Castro, 2025). Other measures of metabolic rate such as resting metabolic rate and maximum metabolic rate are even more limited, hindering large-scale comparative analyses.

One alternative approach to measuring metabolic rate directly is to infer it from morphological characteristics that relate to metabolic rate, dubbed ‘metabolic morphology’ (VanderWright et al., 2020, 2024). Two key metabolic processes, respiration and osmoregulation (Chen et al., 2023; Wegner, 2023), originate at the gills, thus determining our starting point for metabolic morphology. Internally, gills are composed of gill arches with gill filaments covered in lamellae (Bigman et al., 2018; Wegner, 2015). In fishes, gill surface area and metabolic rate are positively interrelated (Bigman et al., 2018; Gillooly et al., 2016; Kuparinen et al., 2022; Scheuffele et al., 2021). Gill surface area also consistently scales positively with body mass within and across species (Bigman et al., 2018; Gillooly et al., 2016; Kuparinen et al., 2022; Scheuffele et al., 2021), and gill surface area is proportionally greater in more active species than less active species (Bigman et al., 2018).

As with metabolic rate, there are challenges to measuring gill surface area. Measuring gill area requires lethal sampling, which is undesirable for threatened species. Additionally, gill area measurements are costly and time-consuming, typically requiring 35+ hours of dissection and microscopy to estimate the gill surface area of a single individual (Bigman et al., 2018; VanderWright et al., 2020). This has led to research into whether measures of external gill morphology are related to gill surface area, such as the operculum in teleosts (bony fishes) or gill slits in elasmobranchs (cartilaginous fishes; Chen et al., 2023; Wegner, 2023). Most focus has been on the outflow morphology (i.e., gill slits in sharks) but the inflow morphology (i.e., spiracle size and shape) has yet to be explored. Gill slit height is proportional in size to gill surface area within and across at least five species of shark (VanderWright et al., 2020), and increases with activity level such that more active species have proportionally larger gill slits (VanderWright et al., 2024). Collectively, this work suggests species with proportionally larger respiratory structures have greater metabolic rates.

Importantly, traits such as spiracle length and gill slit height can be readily measured from specimens in minutes (VanderWright et al., 2020) and does not require mortality or physical destruction of museum specimens. Opportunistic measurements can be taken from living specimens during the course of fieldwork, specimens in fish markets (Mehanna et al., 2025; Osman & Samy-Kamal, 2023), along coastlines during stranding events (Bañón et al., 2026; Tietbohl et al., 2025), as well as in natural history collections (Johnson et al., 2023). Globally, natural history collections house over 1.1 billion objects and fishes are particularly well-represented (Johnson et al., 2023). Natural history specimens are often well-preserved, identified by a taxonomic expert, and have important metadata on collection locality, depth, and date of capture (Singer et al., 2020).

We seek to test whether the positive relationship between shark gill size and activity level (VanderWright et al., 2024) also applies to their sister taxa, the rays (Batoidea). While sharks have only one swimming mode, rays exhibit three swimming styles which may differ in broad activity level: caudal propulsion, pectoral undulation, and pectoral oscillation (Martinez et al., 2016; Rosenberger, 2001). Caudal propulsion involves generating thrust using the muscular tail (Martinez et al., 2016; Rosenberger, 2001), and is seen in species such as guitarfishes and torpedo rays. Pectoral undulation (“wiggling”), is where a wave is propagated backwards along the disc or wing with a wave number >1 (Martinez et al., 2016; Rosenberger, 2001), and is associated with less active, benthic species such as skates and stingrays (Compagno, 1990; Martinez et al., 2016; Rosenberger, 2001; White et al., 2022). Finally, pectoral oscillation (“flapping”) is where a wave is propagated along the disc with a wave number <1 (Martinez et al., 2016; Rosenberger, 2001), and is associated with more active, pelagic species such as devil and eagle rays. All rays possess spiracles which are small openings located immediately behind the eyes for water intake through buccal pumping or branchial ventilation (Ebert et al., 2021; Spieler et al., 2013; Summers & Ferry-Graham, 2001). Beyond their primary role of water intake, spiracle morphology and possible additional functions are poorly understood (Tomita et al., 2012a) with most studies focusing on a single species or genus (Spieler et al., 2013; Summers & Ferry-Graham, 2001; Tomita et al., 2012a).

The few analyses that have examined ray spiracle shape strongly suggest it is adaptive to oxygen uptake capacity, external oxygen environment, and activity levels. Total gill area is positively related to the relative opening of the spiracle (spiracle length * width; Duncan et al., 2015)) in near-term embryos of Amazonian freshwater stingrays (Potamotrygonidae; Duncan et al., 2015). Relative spiracle size has also been used to indicate respiratory performance in potamotrygonids where freshwater stingrays with proportionally larger spiracles are more active than species with proportionally smaller spiracles. Specifically, the Smooth Back River Stingray (*Potamotrygon orbignyi*) is a more active species with proportionally larger spiracles than the Long-Tailed River Stingray (*Plesiotrygon iwamae*; Duncan et al., 2015; Lonardoni et al., 2009). In the Reef Manta (*Mobula alfredi*), spiracles shrink rapidly after birth (Tomita et al., 2012a), possibly within a few days, as neonates shift from buccal pumping of uterine fluid to ram-ventilation while free-swimming in open water (Tomita et al., 2012a).

Here, we take the first step in understanding the metabolic morphology of rays by evaluating the allometry of respiratory morphology. Using measurements of 550 museum specimens from 88 ray species across all four orders, we ask the following four questions: (i) does spiracle length scale with spiracle surface area within and across species (i.e., so spiracle length may be used to approximate spiracle surface area), (ii) does the scaling of spiracle length and gill slit height with body size differ within and across species (do these features grow as a constant proportion with body size), (iii) how are spiracle length and gill slit height related to one another within and across species, and (iv) does the relationship between spiracle length and gill slit height vary by swimming style? We hypothesize: (i) spiracle length will be strongly positively related to spiracle surface area (with an expected slope of ∼2 based on geometric scaling of the relationship between diameter and area of a circle), (ii) spiracle length and gill slit height scale proportionally with body length (proportional size is isometric, i.e. slope =1, as body size increases), (iii) spiracle length and gill slit height will be positively isometrically related to one another, and (iv) the slope of the positive relationship between spiracle length and gill slit height will vary by swimming style, where metabolically active pectoral oscillators will have a steeper slope.

## Materials and Methods

We first describe how morphological data were measured and then describe our statistical analyses.

### Data Collection

We measured 550 rays from 88 species sourced from six natural history collections: Beaty Biodiversity Museum (BBM; Vancouver, Canada), California Academy of Sciences (CAS; San Francisco, United States of America), National Museum of Natural History (MNHN; Paris, France), Natural History Museum (NHMUK; London, United Kingdom), South Africa Institute for Aquatic Biodiversity (SAIAB; Makanda, South Africa), and Zoological Museum Hamburg (ZMH; Hamburg, Germany). We verified specimen identification using field guides (e.g., *Rays of the World*; Last et al., 2016)), taxonomic literature (e.g., Weigmann et al., 2021), and in some cases, consultation with taxonomic experts (D.A. Ebert & S. Weigmann). Specimens that could not be confidently identified to species were excluded from this analysis.

We measured total length (TL; snout tip to posterior edge of the caudal fin, trunk, or tail), disc width (DW; maximum distance between pectoral fin tips), disc length (DL; snout tip to posterior edge of the pectoral fin), and gill slit height (GSH; length of gill opening), following the shape of the gill slit, for all gills slits to the nearest mm (Fig. 1). The height of each gill slit was averaged across all gill slits of each individual (i.e. 10 or 12 gill slits). We also photographed each specimen and imported images into ImageJ to measure spiracle length (SPL; maximum length of the spiracular opening) and spiracle surface area (surface area of the external opening, regardless of whether the spiracle was “open” or “closed”).

**Fig. 1.**
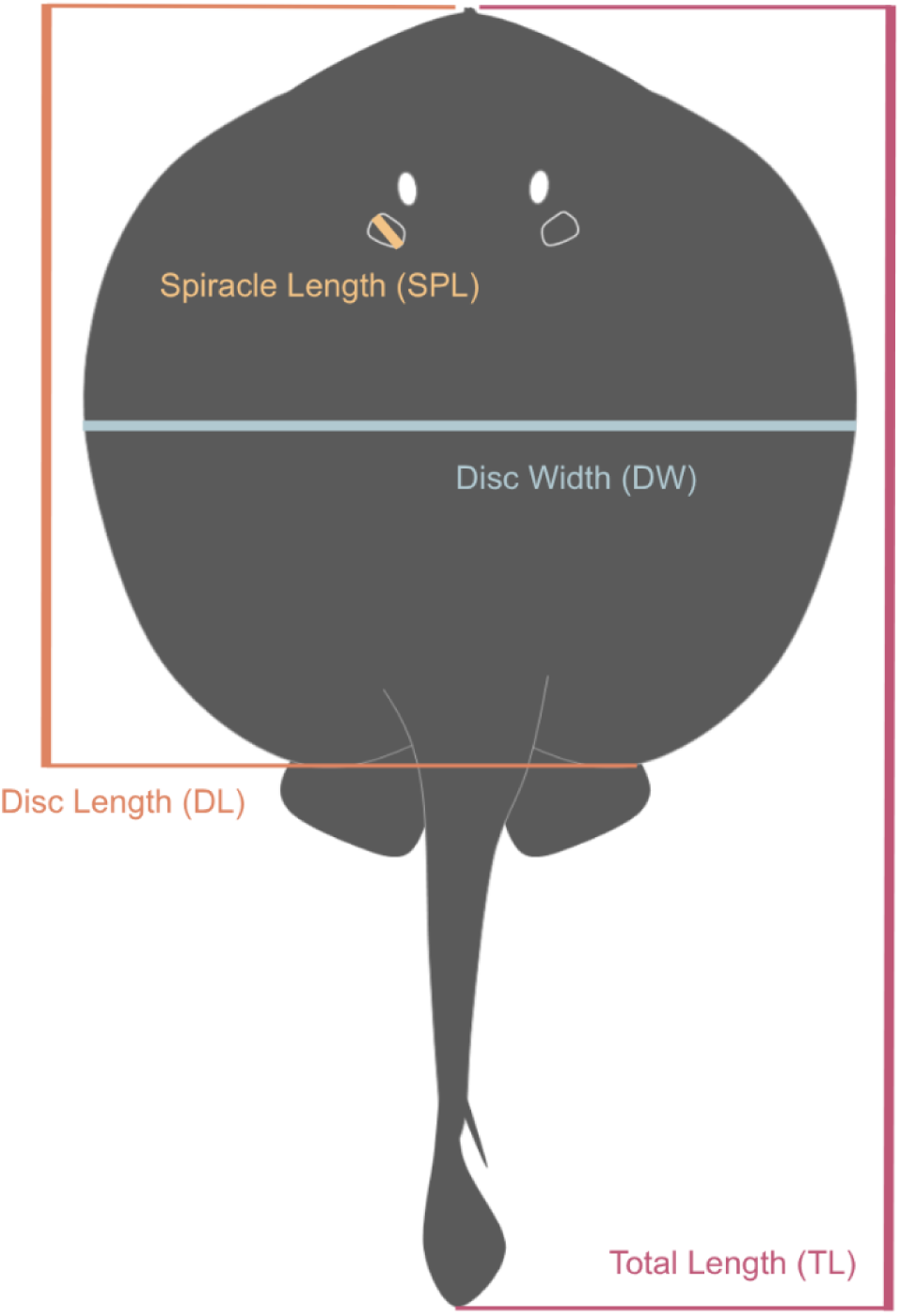
Measurements taken on the dorsal surface of ray specimens. Silhouette of *Urobatis jamaicensis* from phylopic.org (open license, credit to Matt Kolmann) is unchanged except for colouration.

The 88 species spanned all four ray orders (Myliobatiformes, Rajiformes, Rhinopristiformes, and Torpediniformes), and represent 77% (20 of 26) of ray families (Last et al., 2016; Weigmann, 2016). A full species list, including the number of specimens per species, is available in Table S1.

### Statistical Analysis

In order to answer all four of our questions, we fit Bayesian generalized linear multilevel models using the brms package (Bürkner, 2017) with phylogeny as a random effect in R v4.6.1. Prior to analysis, all data were log_10_ transformed to estimate scaling relationships on a linear scale. All models were fit with four chains each of 8,000 iterations with 4,000 warm-up iterations (12,000 iterations total). Model convergence was assessed by ensuring R-hat values were equal to one and effective sample sizes (ESS) were greater than 1,000.

For our first question, whether the scaling of spiracle length and spiracle surface area differs within and across species, our model in R code format was:

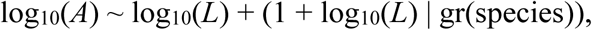

where *A* is spiracle surface area and *L* is spiracle length. This model structure accounts for shared history and heritability of traits while allowing each species to have their own intercept and slope. In taxonomic literature, the term spiracle length is universally used, however, spiracle diameter would be more appropriate as spiracles are more circular to rectangular rather than square. Nevertheless, to avoid confusion, we chose to adhere with taxonomic convention and use the term length. The geometric null expectation based on the relationship between the diameter and area of a circle is A = πd²/4, thus A ∝ d² on a log-log scale, log(A) = log(π/4) + 2·log(d). Therefore a slope of 2 is consistent with isometric growth, where slopes <2 imply spiracles get narrower (elliptical or irregular) and slopes >2 imply spiracles grow wider than long as species grow.

For our second question, whether the allometric scaling of spiracle length and gill slit height with body size differs within and across species, we separated species based on which measurement would best describe body size. We followed the standard used in taxonomic literature where morphometrics are described as proportions of either total length or disc width based on tail morphology. Species with trunk-like tails are described using proportions of total length while species with whip-like tails are described using proportions of disc width. For example, skate (order Rajiformes) and electric ray (order Torpediniformes) measurements are presented as proportions of total length (% TL; Ebert et al., 2015; Iglésias & Lévy-Hartmann, 2012) while eagle ray (families Aetobatidae and Myliobatidae) measurements are presented as proportions of disc width (% DW; Ruocco et al., 2012; White et al., 2016). We then calculated relative spiracle length and relative gill slit height by dividing spiracle length and mean gill slit height by either TL (n = 66 species) or DW (n = 22 species) for each specimen. The structure of our models was:

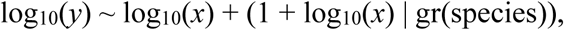

where *y* is relative spiracle length or relative gill slit height and *x* is body size (TL or DW; Table S2).

For our third and fourth questions, how spiracle length and gill slit height are related to one another and whether that relationship varies by swimming style, we assigned swimming style (caudal propulsion, pectoral oscillation, and pectoral undulation) to species based on Compagno (1990) and White et al. (2022). For example, all Rhinopristiformes were classified as caudal propulsion species while all Aetobatidae were classified as pectoral oscillation species. We were able to answer both questions using one model:

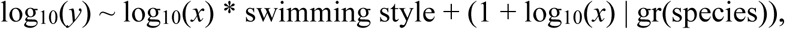

where *y* is relative gill slit height, *x* is spiracle length, and swimming style is a categorical variable with three levels (caudal propulsion, pectoral oscillation, and pectoral undulation).

To account for the shared evolutionary history of traits and non-independence of species, we incorporated phylogenetic trees from Stein et al. (Stein et al., 2018). We adopted recent taxonomic changes (Table S3) and added recently described species missing from the trees using the `phytools` (Revell, 2012) and `ape` (Paradis & Schliep, 2019) packages in R. We added more recently described species (e.g., the Malagasy Blue-Spotted Guitarfish, *Acroteriobatus andysabini*; Weigmann et al., 2021) to the root of the corresponding genus with the `add.species.to.genus` function and assigned branch lengths using the Grafen method, which accounts for shared ancestry without requiring divergence times (Grafen, 1989). We then trimmed tree tips to match the species present in our morphological dataset. Species excluded from analysis and the reason for exclusion are available in Table S4.

All data and code available in GitHub (https://github.com/raitchison/RayAllometry.git).

## Results

We first qualitatively describe the morphology of the spiracles and gills, and then report the results of our statistical models.

### Spiracle and Gill Slit Morphology

Spiracle shape varied widely across species, ranging from quadrilateral (Fig. 2B) to rounded (Fig. 2E) to tear-drop shaped (Fig. 2G). Some, but not all, species in Rhinopristiformes and Torpediniformes possessed papillae (small slender or knob-like protrusions) around the inner edge of the spiracle. For example, the guitarfishes (Rhinobatidae) and giant guitarfishes (Glaucostegidae) possessed 1–2 papillae on the posterior edge of the spiracle (Fig. 2G) while the Bowmouth Guitarfish (Rhinidae) did not (Fig. 2H). Similarly, within the electric torpedo rays (Torpedinidae), species in the genus *Torpedo* possessed papillae around the lateral and posterior edges of the spiracle (Fig. 2I) while species in the genus *Tetronarce* had no papillae.

**Fig. 2.**
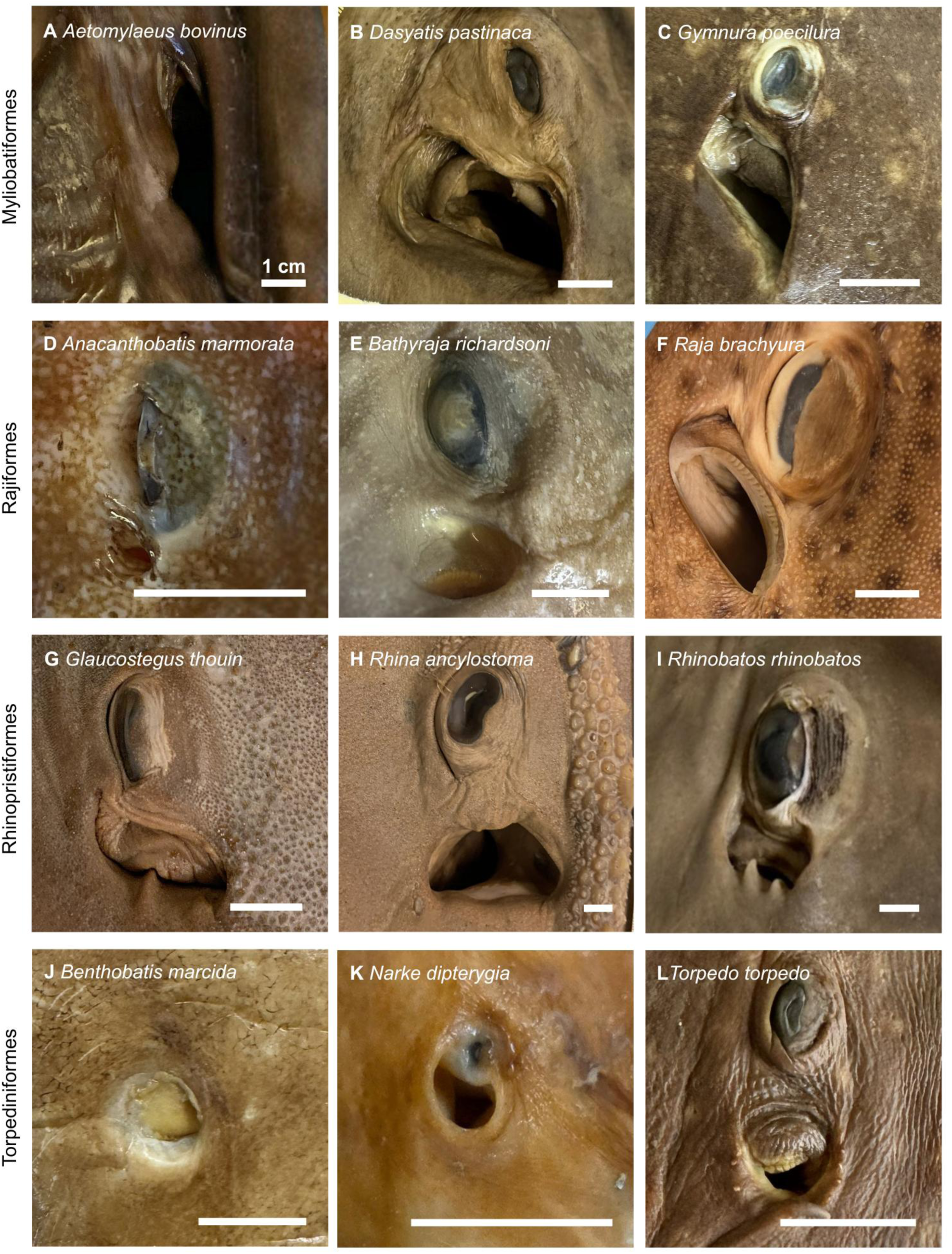
Variation in ray spiracle shape and position relative to the eye. Each row contains three different species from each of the four ray orders: Myliobatiformes (A, B, C, note: *Aetomylaus bovinus* eye is not pictured), Rajiformes (D, E, F), Rhinopristiformes (G, H, I), and Torpediniformes (J, K, L; note: *Benthobatis marcida* eye is greatly reduced). Scale bars = 1 cm.

Gill slit morphology also varied across species, ranging from “wave-shaped” (Fig. 3A) to “C-shaped” (Fig. 3E) to fairly straight (Fig. 3H). The last gill slit (fifth, or sixth for the Sixgill Stingray, *Hexatrygon bickelii*) was often a different shape from the first four gill slits (seen most clearly in Fig. 3F).

**Fig. 3.**
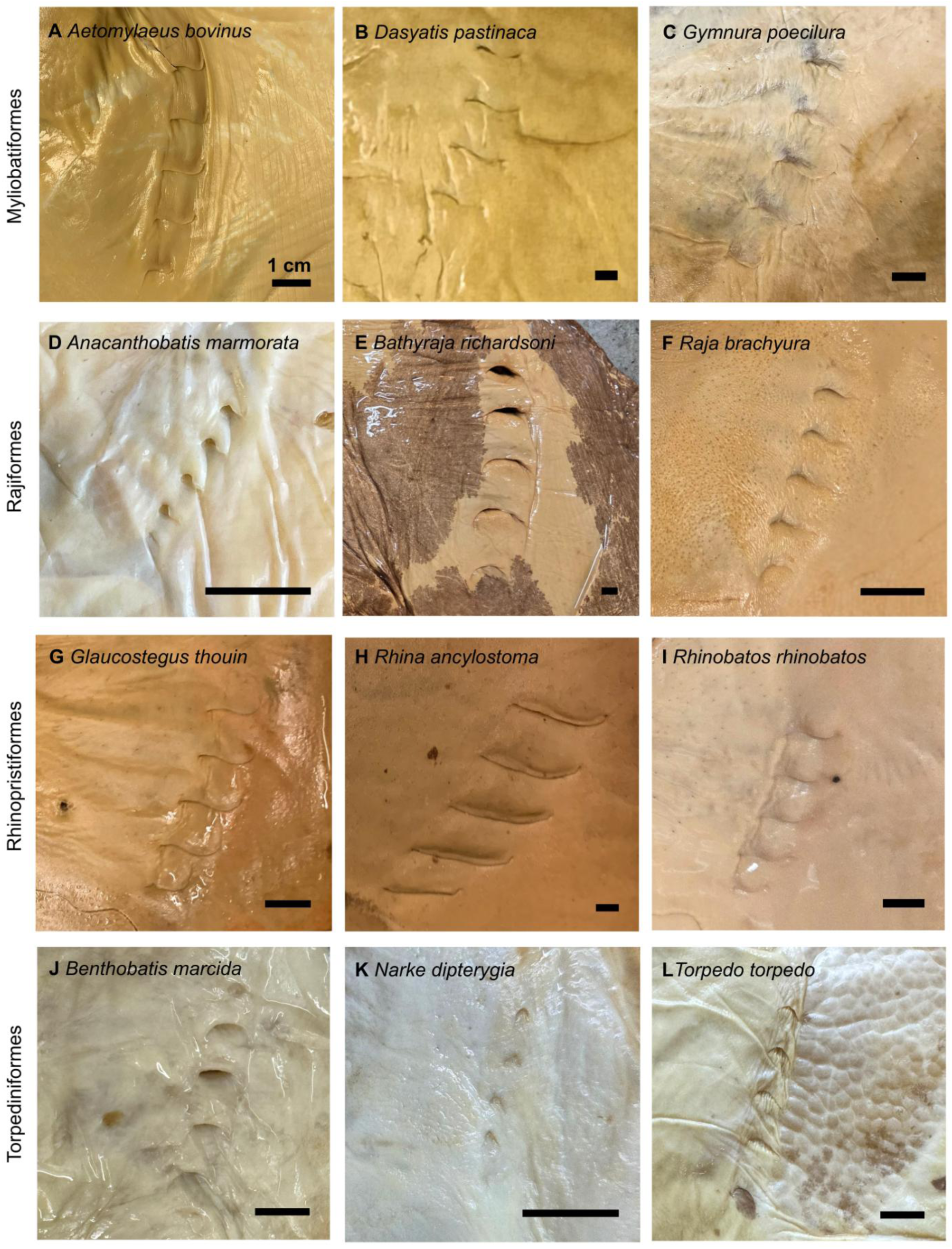
Variation in ray gill slit shape. Each row contains three different species from each of the four ray orders: Myliobatiformes (A, B, C), Rajiformes (D, E, F), Rhinopristiformes (G, H, I), and Torpediniformes (J, K, L). Scale bars = 1 cm.

### Scaling of Spiracle Length with Spiracle Area

Spiracle length and spiracle surface area were positively related such that species with larger spiracle lengths had larger surface areas (Fig. 4). Based on geometric power laws where area is directly proportional to the square of its characteristic linear dimension, the expected coefficient for the relationship between spiracle length and spiracle surface area is 2. In our model, the global (interspecies) slope was 1.94 (95% CI: 1.51 to 2.34) and the credible intervals for the intraspecies slope estimates crossed 2 for most species (n = 63 out of 88 species; Figs 4, S1–S2).

**Fig. 4.**
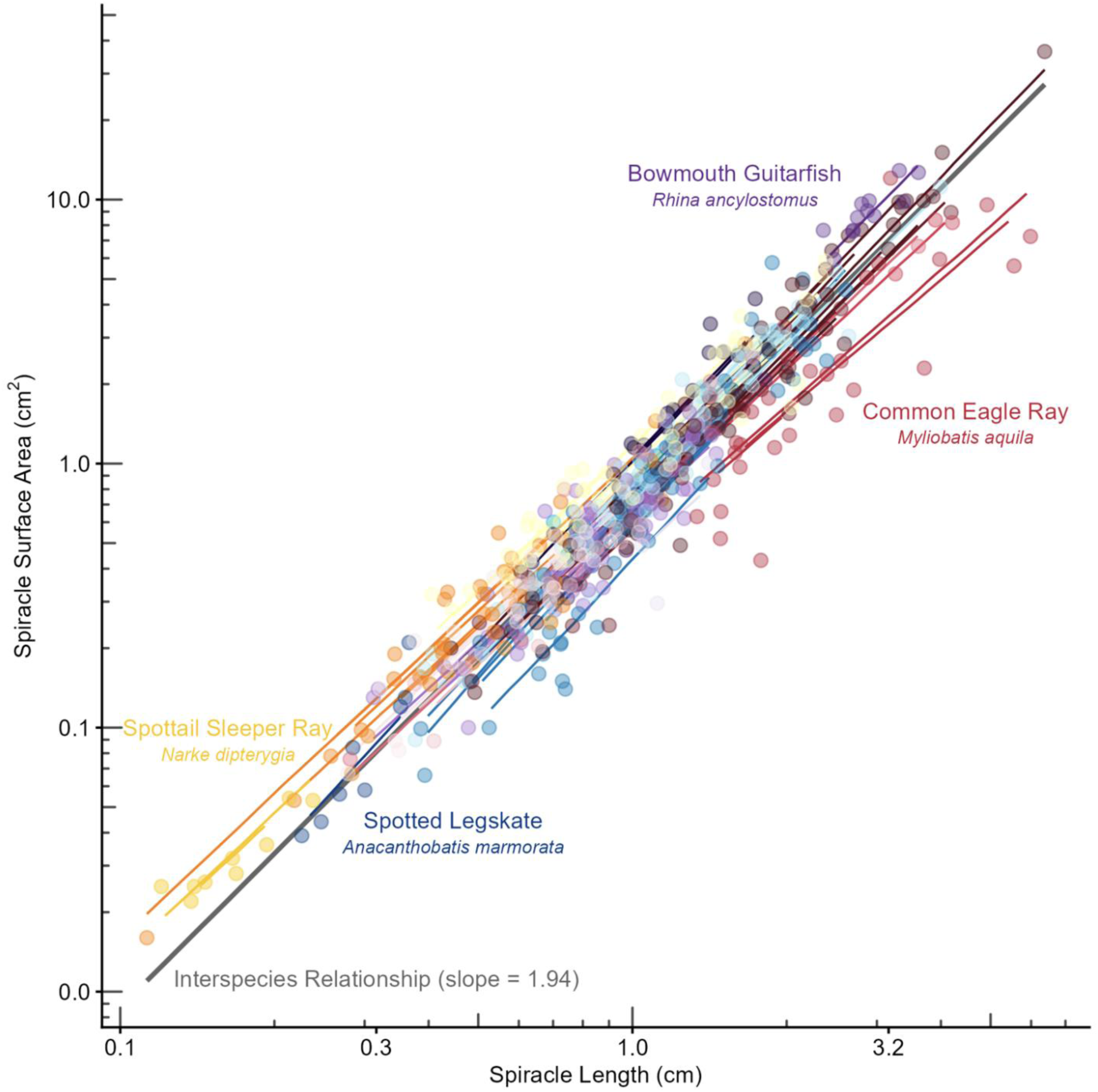
Relationship between spiracle length (cm) and spiracle surface area (cm^2^). Points represent individuals (n = 550) and are colored by family within orders: Myliobatiformes (reds and pinks), Rajiformes (blues), Rhinopristiformes (purples), and Torpediniformes (oranges and yellows). Each line represents the species-specific relationship from the model (see also Figs S1–S2). The grey line represents the interspecies (global) slope from the model.

Of the 25 species whose slopes varied significantly from 2 (i.e., the 95% CI did not reach 2), twenty-three species exhibited slope estimates <2 which means the spiracle surface area is smaller than expected based on spiracle length. Only two species slopes were >2 meaning the spiracle surface area is greater than expected based on spiracle length; Figs S1–S2). The shallowest modeled slope was in the Common Eagle Ray (*Myliobatis aquila*; 1.67, 95% CI: 1.47 to 1.86) which has a narrow rhomboid shape, and the steepest modeled slope was in Richardson’s Ray (*Bathyraja richardsoni* 2.21, 95% CI: 2.04–2.42; Figs 2E–3E) which has a squat ovoid shape.

### Isometry of Spiracle Length and Gill Slit Height

We found that relative spiracle length and gill slit height were a nearly consistent proportion of body size (total length or disc width) for most species (n = 75 and n = 63 out of 88 species for spiracle length and gill slit height, respectively; Figs 5–6, S3–S6). In other words, the ratio of spiracle length and gill slit height to body size generally does not change as individuals grow in body size through ontogeny.

**Fig. 5.**
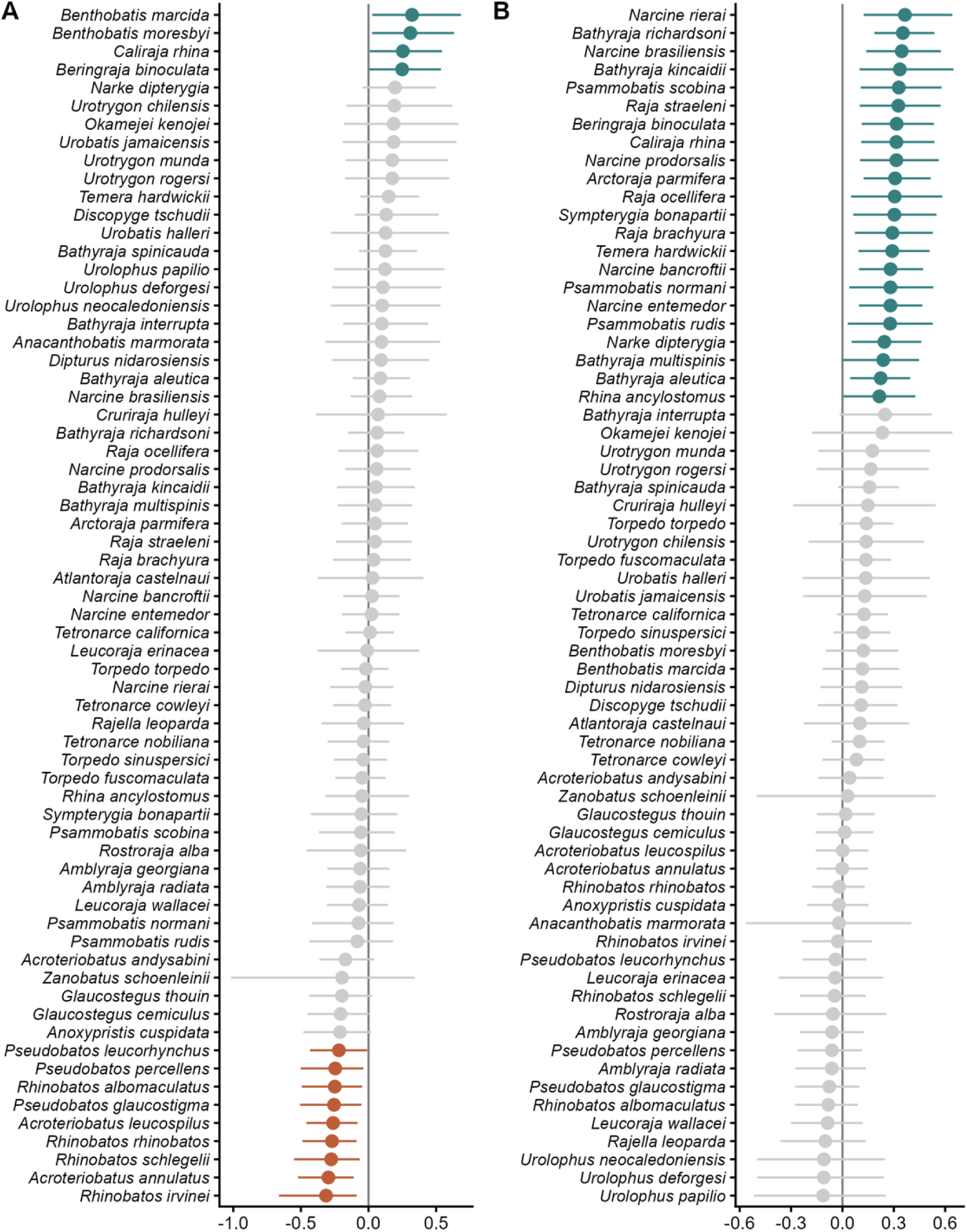
Coefficient plots of scaling relationships for species defined by total length (n = 88). (A) relative spiracle length, (B) relative gill slit height. Lines for each coefficient indicate the 95% credible interval. Color indicates the statistical significance based on the confidence interval: above 0 (turquoise), not different from 0 (grey), and below 0 (rust). See also Figs S3–S4.

**Fig. 6.**
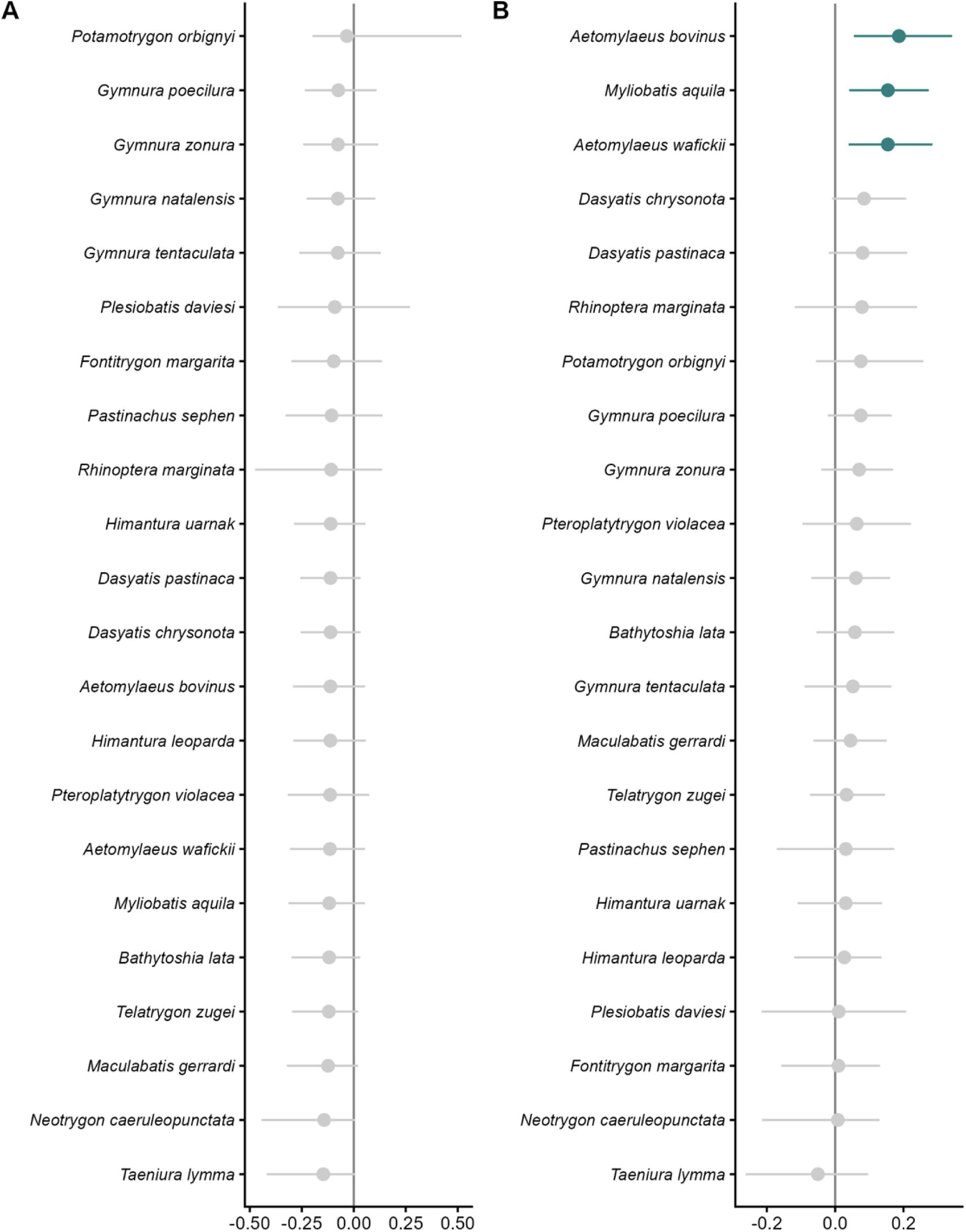
Coefficient plots of scaling relationships for species defined by disc width (n = 22). (A) relative spiracle length, (B) relative gill slit height. Lines for each coefficient indicate the 95% credible interval. Color indicates the statistical significance based on the confidence interval: above 0 (turquoise), not different from 0 (grey), and below 0 (rust). See also Figs S5–S6.

For species defined by total length, mean relative spiracle length varied from 1.53% TL in the Spottail Sleeper Ray (*Narke dipterygia*) to 3.99% TL in the Striped Panray (*Zanobatus schoenleinii*) and mean gill slit height varied from 0.93% TL in the Spotted Legskate (*Anacanthobatis marmorata*) to 2.93% TL in the Pacific Electric Ray (*Tetronarce californica*). We found that both spiracle length and gill slight height grow isometrically with total length. The global (interspecific) slopes were isometric and indistinguishable from 0 (relative spiracle length *β* = 0.03, 95% CI -0.69 to 0.74; and relative gill slit height *β* = 0.10, 95% CI -0.51 to 0.70; Figs 5, S3–S4). For spiracle length, four species exhibited slope estimates greater than 0, meaning spiracles grow proportionally larger with increasing body size, and nine species slopes were less than 0, meaning spiracles grow proportionally smaller with increasing body size (Figs 5, S3). For relative gill slit height, 22 species had slope estimates greater than 0, meaning gill slits grow proportionally larger with increasing body size (Figs 5, S4).

For species in which maximum size is defined by disc width, mean relative spiracle length varied from 3.00% DW in the Zonetail Butterfly Ray (*Gymnura zonura*) to 8.29% DW in Bluespotted Ribbontail Ray (*Taeniura lymma*) and mean gill slit height varied from 1.62% DW in the Bull Ray (*Aetomylaeus bovinus*) to 3.97% DW in the Bluespotted Ribbontail Ray (*Taeniura lymma*). We found that both spiracle length and gill slight height grow isometrically with disc width. The global (interspecific) slopes were isometric and indistinguishable from 0 (relative spiracle length *β =* -0.08, 95% CI: -0.32 to 0.20; and relative gill slit height *β =* 0.05, 95% CI: -0.30 to 0.27; Figs 6, S5–S6). For relative spiracle length, no species exhibited slope estimates greater than or less than 0 (Figs 6, S5). For relative gill slit height, three species had slope estimates greater than 0, meaning gills grow proportionally larger with increasing body size (Figs 6, S6).

### Scaling of Spiracle Length and Gil Slit Height

Spiracle length and mean gill slit height were positively related across all three swimming styles (caudal propulsion, pectoral oscillation, pectoral undulation) and the slope of each relationship was near 1 (Fig. 7, Table 1). Slopes varied from 0.84 (95% CI: 0.26 to 1.42) in pectoral undulating species to 1.21 (95% CI: 0.46 to 2.04) in pectoral oscillating species. Although the slopes are not statistically different from each other, as indicated by their overlapping credible intervals (Fig. 7, Table 1), pectoral undulation and caudal propulsion are most similar to each other (0.84 and 0.85, respectively) and different from pectoral oscillation (1.21).

**Fig. 7.**
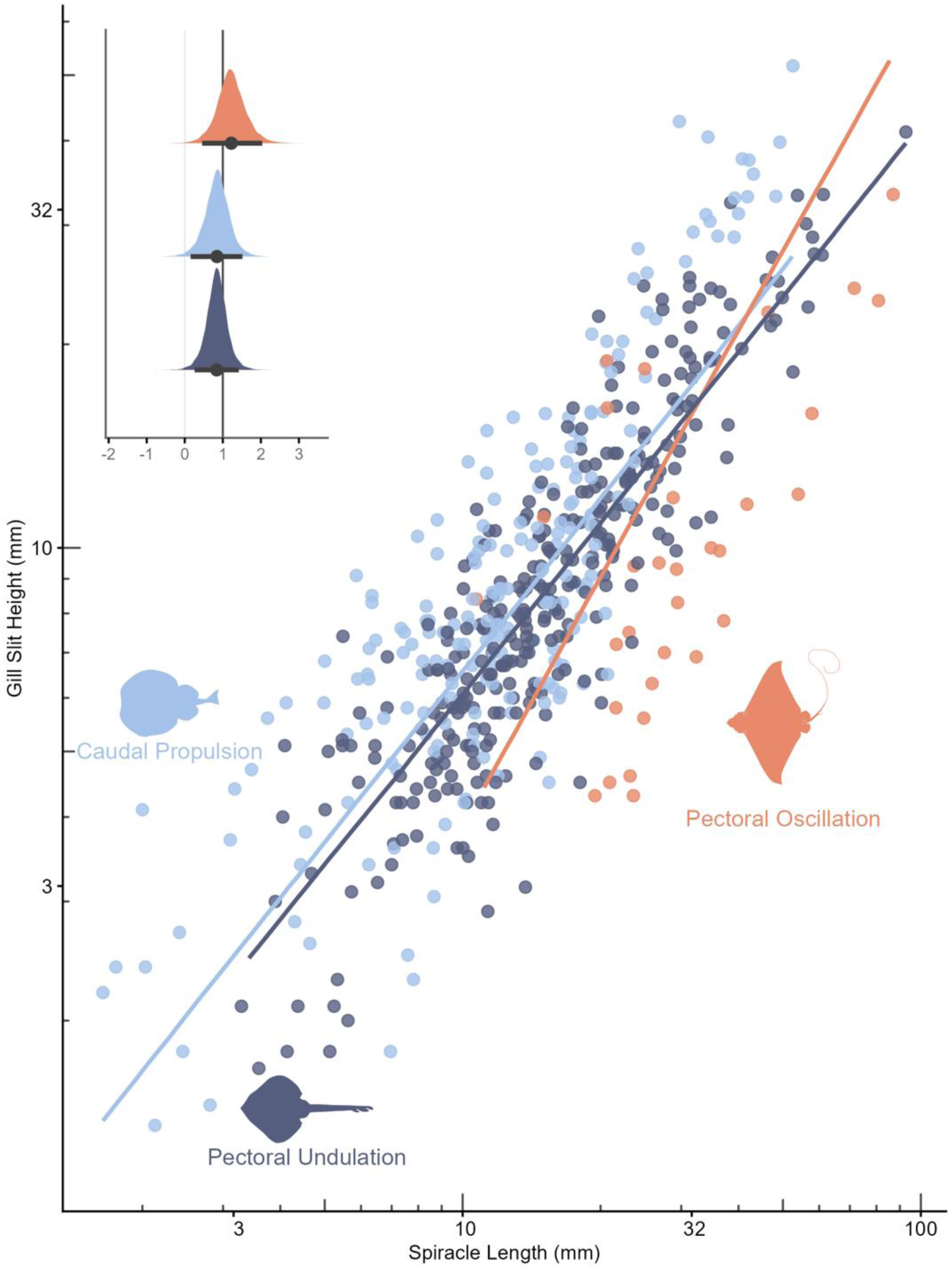
Relationship between spiracle length and gill slit height for each swimming style. Color indicates swimming style: coral (pectoral oscillation), blue (caudal propulsion), and slate (pectoral undulation). Inset shows model coefficient estimates and posterior probabilities of spiracle length and mean gill slit height for each swimming style, and lines for each coefficient indicate the 95% credible interval.Silhouettes from phylopic.org (open license, credit to Matt Kolmann, Steven Traver, and Guillaume Dera) are unchanged except for orientation and colouration.

**Table 1.** Model summary table for the relationship between spiracle length and gill slit height of each swimming style.

| Swimming Style | Body Measurement | Coefficient | 95% CI | Number of species | Number of individuals |
| --- | --- | --- | --- | --- | --- |
| **Pectoral Oscillation** | Disc Width (DW) | 1.21 | 0.46–2.04 | 4 | 31 |
| **Caudal Propulsion** | Total Length (TL) | 0.85 | 0.15–1.52 | 31 | 204 |
| **Pectoral Undulation** | Disc Width (DW) or Total Length (TL) | 0.84 | 0.26–1.42 | 53 | 315 |

## Discussion

Overall, we found strong positive interrelationships between respiratory structures. Specifically, spiracle length and gill slit height are closely related within and across species, such that species with larger spiracles have longer gill slits – which is broadly consistent with greater inflow area related to greater outflow area. This is also consistent with the overarching idea that species with larger spiracles and longer gill slits likely have greater metabolic rate, aerobic scope, and activity levels (Bigman et al., 2018; Iliou et al., 2023; VanderWright et al., 2024). Reviewing our own results alongside three complementary studies, we suggest that this tight integration of respiratory morphology arises early in life and extends throughout ontogeny with spiracle length and gill slit height growing proportionally (isometrically) with body size. Specifically, we consider (i) whether respiratory morphology varies with more detailed measures of activity, (ii) metabolic hypotheses based on the biogeographic distribution of water temperature that might reveal the adaptive nature of spiracle and gill slit size, and (iii) specific hypotheses on the change in spiracle size upon parturition for species with different reproductive modes, before (iv) concluding with three recommendations for evaluating the utility of and expanding the use of respiratory morphological knowledge.

Our results further suggest that more active species have larger metabolic morphologies. Specifically, pectoral oscillation species, which are pelagic, tend to have larger respiratory morphologies than the other two swimming styles (caudal propulsion and pectoral undulation), which are mostly benthic. In addition, our findings suggest either spiracle length or gill slit height could be used to evaluate activity levels since spiracle length and gill slit height scale proportional to each other (Fig. 7, Table 1). Defining activity levels, however, presents an additional challenge. In fishes, activity level has been parameterised with the aspect ratio of the caudal fin (Palomares & Pauly, 1989; Sambilay, 1990; VanderWright et al., 2024). The mechanistic link between caudal fin shape and activity was confirmed by the strong positive relationship between Caudal Fin Aspect Ratio and swimming speeds where sharks with homocercal or lunate tails swim faster than species with heterocercal or paddle-like tails (Iliou et al., 2023). A mechanistic link between ray fin shape and activity has not yet been established, and presents a challenge since rays are morphologically diverse (i.e. not all species have caudal fins) and exhibit three swimming styles (Compagno, 1990; Martinez et al., 2016; White et al., 2022). We also caution that our analysis only includes four pectoral oscillation species (Table 1).

Our work suggests three main avenues of future research testing the adaptive nature of respiratory morphology. The first is relating respiratory morphology to more detailed measures of activity. In sharks, gill slit height is greater in larger and shallower species (and endothermic species) have higher metabolic throughput as indexed by gill slit height (oxygen uptake) and caudal fin aspect ratio (oxygen expenditure; VanderWright et al., 2024). Further, more active species (indexed by caudal fin aspect ratio) have faster average swimming speeds in the wild as measured by telemetry (Iliou et al., 2023). Based on what we have learned from sharks, we hypothesize that ray respiratory morphology will relate to ray swimming speeds and measures of genetic connectivity, such as isolation by distance (Kipperman et al., 2025).

The second is the exploration of respiratory morphology across different environmental temperatures and oxygen levels. Across all our analyses, a large proportion of the species that did not follow the general pattern observed were skates (Rajiformes). Of the 63 species of skate present in our dataset, 14 had statistically significantly different scaling relationships between relative mean gill slit height and total length. Skates are primarily benthic (Compagno, 1990; White et al., 2022) and exhibit ontogenetic shifts in depth, and thus oxygen availability, that are often inverse with body size. For example, smaller individuals of the Flapper Skate, *Dipturus intermedius*, and the Shagreen Ray, *Leucoraja fullonica*, are found at deeper depths than larger individuals (Bisch et al., 2022; Thorburn et al., 2021). Interestingly, species in our dataset from the same two genera (*Dipturus* and *Leucoraja*) did not have a statistically significant difference in relative mean gill slit height across body sizes. However, a large proportion of these 14 species are found at high latitudes, which have documented latitudinal gradients in life history traits such as size-at-maturity (Frisk & Miller, 2009; Winton et al., 2013). Geographic location, including latitude, were not factored into our analyses and the individuals we measured cover a range of latitudes and may, as a result, cover a range of metabolic morphologies. Thus, additional work is needed to investigate whether latitude, and associated gradients in temperature and oxygen, influence metabolic morphology within species.

The third research direction we suggest based on our work is a wider evaluation of changes to spiracle and gill morphology across all life states based on reproductive strategy. We highlight that while our analysis covers a range of body sizes within each species, it does not cover the full range of potential body sizes (particularly neonates or young-of-the-year) or consider reproductive strategy. Based on changes in spiracle size of the Reef Manta (*Mobula alfredi*) upon birth (Tomita et al., 2012b), we predict marked transitions in spiracle size in the ram-ventilating matrotrophic species (other devil rays, eagle rays, and cownose rays), but not in buccal-pumping matrotrophic species (stingrays). We also expect lecithotrophic neonates (guitarfishes and torpedo rays) to have proportionally larger spiracles prior to birth compared to oviparous neonates (skates) which should have no difference in proportional spiracle size before and after hatching.

While our work spans some taxonomic breadth, we acknowledge that our analysis includes a fraction of ray species diversity (88 species, ∼14% of 630+ species; Last et al., 2016; Weigmann, 2016). Thus, we have three recommendations to expand species sampling to support wider hypothesis testing. In addition to museum specimens, additional data could be sourced from taxonomic literature which includes some descriptions of the spiracles and gills. We caution, however, that these descriptions are often sparse. While spiracle length is often reported, gill slit heights are reported inconsistently across orders. Specifically, only the third gill slit height is reported for guitarfishes (Weigmann et al., 2021), while the first, third, and fifth gill slit heights are reported for electric rays (Ebert et al., 2015). First, we recommend taxonomists report all gill slit heights. Second, we recommend the next edition of Rays of the World incorporates taxonomically accurate ventral illustrations. This would allow extraction of gill slit height directly from these images, as has been successful in sharks (VanderWright et al., 2020, 2024). Third, we encourage taxonomists to continue exploring spiracle morphology as a diagnostic character for distinguishing species and encourage scientific illustrators to continue working with taxonomists to ensure spiracles are drawn accurately. Spiracle morphology is used to distinguish some genera (e.g. *Tetronarce* and *Torpedo* within Torpedinidae) and species. For example, the Panther Torpedo (*Torpedo panthera*) and Persian Gulf Torpedo (*T. sinuspersici*) are morphologically similar and have overlapping geographic ranges, however, the Panther Torpedo has seven papillae around the spiracle while the Persian Gulf Torpedo has 9–10 papillae around the spiracle (Khan & Osmany, 2021; Last et al., 2016). Although currently used primarily as a diagnostic tool, we wonder whether spiracle papillae vary more by environment (e.g. more papillae prevent sand from entering the spiracle) than taxonomy, thus raising questions about the function of spiracle papillae.

In conclusion, our comparative analysis of the intra- and interspecific allometry of respiratory morphology offers broad evidence for the adaptive nature of respiratory morphology of rays. This complements similar findings in sharks that show that gill slit morphology is tightly related to metabolic rates and activity levels. Given the challenge of measuring metabolic rates for large free-swimming animals, this work offers the promise of a metabolically informed understanding of life history traits, population growth, extinction risk, and ecological function.

## Acknowledgements

The authors would like to sincerely thank collections staff for their generous support and assistance, especially Nicolas Bailly (BBM), Eric Taylor (BBM), David Catania (CAS), Zouhaira Gabsi (MNHN), Jonathon Pfliger (MNHN), Patrice Pruvost (MNHN), Rupert Collins (NHMUK), Nkosinathi Mazungula (SAIAB), Timo Moritz (ZMH), and Thilo Weddehage (ZMH). For their help measuring specimens and taking photographs, the authors would also like to thank Jonathan Eged and Sarah Gravel. Finally, the authors would like to thank Dave Ebert and Simon Weigmann for their taxonomic expertise and assistance in identifying museum specimens.

## References

Bañón, R., de Carlos, A., Pérez, A. E., Comesaña, Á. S., & Falcón, J. M. (2026). New records and evidence for the Atlantic distribution of Bregmaceros atlanticus (Gadiformes: Bregmacerotidae). Marine Biodiversity, 56(4), 69. 10.1007/s12526-026-01679-5

Bigman, J. S., M’Gonigle, L. K., Wegner, N. C., & Dulvy, N. K. (2021). Respiratory capacity is twice as important as temperature in explaining patterns of metabolic rate across the vertebrate tree of life. Science Advances, 7(19), eabe5163. 10.1126/sciadv.abe5163

Bigman, J. S., Pardo, S. A., Prinzing, T. S., Dando, M., Wegner, N. C., & Dulvy, N. K. (2018). Ecological lifestyles and the scaling of shark gill surface area. Journal of Morphology, 279(12), 1716–1724. 10.1002/jmor.20879

Bisch, A., Elliott, S. A. M., Carpentier, A., & Acou, A. (2022). Modelling the distribution of vulnerable skate from fisheries dependent data using imperfect detection. Progress in Oceanography, 206, 102859. 10.1016/j.pocean.2022.102859

Brandl, S. J., Lefcheck, J. S., Bates, A. E., Rasher, D. B., & Norin, T. (2023). Can metabolic traits explain animal community assembly and functioning? Biological Reviews, 98(1), 1–18. 10.1111/brv.12892

Brown, J. H., Gillooly, J. F., Allen, A. P., Savage, V. M., & West, G. B. (2004). TOWARD A METABOLIC THEORY OF ECOLOGY. Ecology, 85(7), 1771–1789. 10.1890/03-9000

Bürkner, P.-C. (2017). brms: An R Package for Bayesian Multilevel Models Using Stan. Journal of Statistical Software, 80. 10.18637/jss.v080.i01

Chen, X., Liu, S., Ding, Q., Teame, T., Yang, Y., Ran, C., Zhang, Z., & Zhou, Z. (2023). Research advances in the structure, function, and regulation of the gill barrier in teleost fish. Water Biology and Security, 2(2), 100139. 10.1016/j.watbs.2023.100139

Compagno, L. J. V. (1990). Alternative life-history styles of cartilaginous fishes in time and space. Environmental Biology of Fishes, 28(1), 33–75. 10.1007/BF00751027

de Castro, F. (2025). FmrBT: A comprehensive data set of field metabolic rates, body mass and ambient temperature. Scientific Data, 12(1), 1589. 10.1038/s41597-025-05868-y

Duncan, W. P., Silva, M. I. da, & Fernandes, M. N. (2015). Gill dimensions in near-term embryos of Amazonian freshwater stingrays (Elasmobranchii: Potamotrygonidae) and their relationship to the lifestyle and habitat of neonatal pups. Neotropical Ichthyology, 13, 123–136. 10.1590/1982-0224-20140132

Ebert, D. A., Dando, M., & Fowler, S. (2021). Sharks of the World: A Complete Guide (2nd edn). Wild Nature Press.

Ebert, D. A., Haas, D. L., & De Carvalho, M. R. (2015). Tetronarce cowleyi, sp. nov., a new species of electric ray from southern Africa (Chondrichthyes: Torpediniformes: Torpedinidae). Zootaxa, 3936(2), 237. 10.11646/zootaxa.3936.2.4

Frisk, M. G., & Miller, T. J. (2009). Maturation of Little Skate and Winter Skate in the Western Atlantic from Cape Hatteras to Georges Bank. Marine and Coastal Fisheries, 1(1), 1–11. 10.1577/C08-014.1

Gillooly, J., Gómez, J., Mavrodiev, E., Rong, Y., & Mclamore, E. (2016). Body mass scaling of passive oxygen diffusion in endotherms and ectotherms. Proceedings of the National Academy of Sciences, 113, 5340–5345. 10.1073/pnas.1519617113

Grafen, A. (1989). The phylogenetic regression. Philosophical Transactions of the Royal Society of London. B, Biological Sciences, 326(1233), 119–157. 10.1098/rstb.1989.0106

Iglésias, S., & Lévy-Hartmann, L. (2012). Bathyraja leucomelanos, a new species of softnose skate (Chondrichthyes: Arhynchobatidae) from New Caledonia. Ichthyological Research, 59, 38–48. 10.1007/s10228-011-0254-y

Iliou, A. S., Vanderwright, W., Harding, L., Jacoby, D. M. P., Payne, N. L., & Dulvy, N. K. (2023). Tail shape and the swimming speed of sharks. Royal Society Open Science, 10(10), 231127. 10.1098/rsos.231127

Johnson, K., Owens, I., Abramov, A., Ametrano, S., Baker, G., Bartolozzi, L., Basseches, J., Beckel, M., Bennett, S., Benvenuti, M., Bettison-Varga, L., Bird, T., Carrasquero, S., Chaney, H., Chernetsov, N., Chiappe, L., Cibois, A., Coles, A., Cooper, S., & Zardoya, R. (2023). A global approach for natural history museum collections. Science, 379, 1192–1194.

Khan, M., & Osmany, H. (2021). SPECIES COMPOSITION AND DISTRIBUTION OF ELECTRIC RAYS (CLASS: PISCES; SUBCLASS: ELASMOBRANCHII; ORDER: TORPEDINIFORMES) FROM PAKISTAN. 725–743.

Kipperman, M., Jabado, R., Haque, A., Fernando, D., Anjani, P. A. D. L., Spaet, J., & Humble, E. (2025). Population structure and genetic diversity of the Critically Endangered bowmouth guitarfish (Rhina ancylostomus) in the Northwest Indian Ocean. Peer Community Journal, 5. 10.24072/pcjournal.623

Kleiber, M. (1947). BODY SIZE AND METABOLIC RATE. Physiological Reviews, 27(4), 511–541. 10.1152/physrev.1947.27.4.511

Kuparinen, A., Gielewski, D., & Hutchings, J. A. (2022). Gill area explains deviations from body size–metabolic rate relationship in teleost fishes. Journal of Fish Biology, 101(1), 308–311. 10.1111/jfb.15084

Last, P. R., White, W. T., De Carvalho, M. R., Séret, B., Stehmann, M. F. W., & Naylor, G. J. P. (2016). Rays of the World. CSIRO Publishing.

Lonardoni, A., Oliveira, E., & Goulart, E. (2009). Trophic ecomorphology of Potamotrygon falkneri and Potamotrygon motoro (Chondrichthyes—Potamotrygonidae) on the upper Paraná river floodplain, Brazil. Pan-American Journal of Aquatic Sciences, 4, 436–445.

Martinez, C. M., Rohlf, F. J., & Frisk, M. G. (2016). Re-evaluation of batoid pectoral morphology reveals novel patterns of diversity among major lineages: PECTORAL DIVERSITY OF BATOID FISHES. Journal of Morphology, 277(4), 482–493. 10.1002/jmor.20513

Mehanna, S. F., Samy-Kamal, M., Desouky, M. G., El-Mahdy, S. M., Osman, Y. A., Koleib, Z. M., & Ibrahim, R. (2025). Sharks in the Southeastern Mediterranean: Species diversity, fisheries, market, and conservation measures. Regional Studies in Marine Science, 89, 104394. 10.1016/j.rsma.2025.104394

Osman, Y. A., & Samy-Kamal, M. (2023). Diversity and characteristics of commercial Red Sea fish species based on fish market survey: Informing management to reduce the risk of overfishing. Journal of Fish Biology, 102(4), 936–951. 10.1111/jfb.15339

Palomares, M. L. D., & Pauly, D. (1989). A multiple regression model for prediction the food consumption of Marine Fish populations. Marine and Freshwater Research, 40, 259–273. 10.1071/MF9890259

Paradis, E., & Schliep, K. (2019). ape 5.0: An environment for modern phylogenetics and evolutionary analyses in R. Bioinformatics, 35(3), 526–528. 10.1093/bioinformatics/bty633

Pettersen, A. K., White, C. R., & Marshall, D. J. (2016). Metabolic rate covaries with fitness and the pace of the life history in the field. Proceedings of the Royal Society B: Biological Sciences, 283(1831), 20160323. 10.1098/rspb.2016.0323

Revell, L. J. (2012). phytools: An R package for phylogenetic comparative biology (and other things). Methods in Ecology and Evolution, 3(2), 217–223. 10.1111/j.2041-210X.2011.00169.x

Ricklefs, R. E., & Wikelski, M. (2002). The physiology/life-history nexus. Trends in Ecology & Evolution, 17(10), 462–468. 10.1016/S0169-5347(02)02578-8

Rosenberger, L. (2001). Pectoral Fin Locomotion in Batoid Fishes: Undulation Versus Oscillation. Journal of Experimental Biology, 204, 379–394. 10.1242/jeb.204.2.379

Ruocco, N., Lucifora, L., Diaz de Astarloa, J., Mabragaña, E., & Delpiani, M. (2012). Morphology and DNA Barcoding Reveal a New Species of Eagle Ray from the Southwestern Atlantic: Myliobatis ridens sp. nov. (Chondrichthyes: Myliobatiformes: Myliobatidae). Zoological Studies, 51, 862–873.

Sambilay, V. C. (1990). Interrelationships between swimming speed, caudal fin aspect ratio and body length of fishes. Fishbyte, 8(3), 16–20.

Scheuffele, H., Jutfelt, F., & Clark, T. (2021). Investigating the gill-oxygen limitation hypothesis in fishes: Intraspecific scaling relationships of metabolic rate and gill surface area. Conservation Physiology, 9. 10.1093/conphys/coab040

Singer, R. A., Ellis, S., & Page, L. M. (2020). Awareness and use of biodiversity collections by fish biologists. Journal of Fish Biology, 96(2), 297–306. 10.1111/jfb.14167

Spieler, R. E., Fahy, D. P., Sherman, R. L., Sulikowski, J. A., & Quinn, T. P. (2013). The Yellow Stingray, Urobatis jamaicensis (Chondrichthyes: Urotrygonidae): a synoptic review. Caribbean Journal of Science, 47(1), 67–97. 10.18475/cjos.v47i1.a8

Stein, R. W., Mull, C. G., Kuhn, T. S., Aschliman, N. C., Davidson, L. N. K., Joy, J. B., Smith, G. J., Dulvy, N. K., & Mooers, A. O. (2018). Global priorities for conserving the evolutionary history of sharks, rays and chimaeras. Nature Ecology & Evolution, 2(2), 288–298. 10.1038/s41559-017-0448-4

Summers, A. P., & Ferry-Graham, L. A. (2001). Ventilatory Modes And Mechanics Of The Hedgehog Skate (Leucoraja Erinacea): Testing The Continuous Flow Model. Journal of Experimental Biology, 204(9), 1577–1587. 10.1242/jeb.204.9.1577

Thorburn, J., Wright, P. J., Lavender, E., Dodd, J., Neat, F., Martin, J. G. A., Lynam, C., & James, M. (2021). Seasonal and Ontogenetic Variation in Depth Use by a Critically Endangered Benthic Elasmobranch and Its Implications for Spatial Management. Frontiers in Marine Science, 8. 10.3389/fmars.2021.656368

Tietbohl, M. D., Genevier, L. G. C., Krieger, E. C., Kattan, A., Wang, Y., Gokul, E. A., Rodriguez Bravo, L. M., Palm, L., Mele, G., Hoteit, I., & Johnson, M. D. (2025). Extreme marine heatwave linked to mass fish kill in the Red Sea. Science of The Total Environment, 975, 179073. 10.1016/j.scitotenv.2025.179073

Tomita, T., Toda, M., Ueda, K., Uchida, S., & Nakaya, K. (2012a). Live-bearing manta ray: How the embryo acquires oxygen without placenta and umbilical cord. Biology Letters, 8(5), 721–724. 10.1098/rsbl.2012.0288

Tomita, T., Toda, M., Ueda, K., Uchida, S., & Nakaya, K. (2012b). Live-bearing manta ray: How the embryo acquires oxygen without placenta and umbilical cord. Biology Letters, 8(5), 721–724. 10.1098/rsbl.2012.0288

VanderWright, W. J., Bigman, J. S., Elcombe, C. F., & Dulvy, N. K. (2020). Gill slits provide a window into the respiratory physiology of sharks. Conservation Physiology, 8(1), coaa102. 10.1093/conphys/coaa102

VanderWright, W. J., Bigman, J. S., Iliou, A. S., & Dulvy, N. K. (2024). Ecological lifestyle and gill slit height across sharks. Royal Society Open Science, 11(5), 231867. 10.1098/rsos.231867

Wegner, N. C. (2015). Elasmobranch Gill Structure. In Fish Physiology (Vol. 34, pp. 101–151). Academic Press. 10.1016/B978-0-12-801289-5.00003-1

Wegner, N. C. (2023). Chapter 1 - The fish gill: Where fish physiology begins. In D. J. Randall, A. P. Farrell, C. J. Brauner, & E. J. Eliason (Eds), The 50th Anniversary Issue of Fish Physiology: Physiological Systems and Development (Vol. 40, pp. 1–7). Academic Press. 10.1016/bs.fp.2023.09.003

Weigmann, S. (2016). Annotated checklist of the living sharks, batoids and chimaeras (Chondrichthyes) of the world, with a focus on biogeographical diversity: Annotated global checklist of chondrichthyes. Journal of Fish Biology, 88(3), 837–1037. 10.1111/jfb.12874

Weigmann, S., Ebert, D. A., & Séret, B. (2021). Resolution of the Acroteriobatus leucospilus species complex, with a redescription of A. leucospilus (Norman, 1926) and descriptions of two new western Indian Ocean species of Acroteriobatus (Rhinopristiformes, Rhinobatidae). Marine Biodiversity, 51(4), 58. 10.1007/s12526-021-01208-6

White, W. T., Last, P. R., & Baje, L. (2016). Aetomylaeus caeruleofasciatus, a new species of eagle ray (Myliobatiformes: Myliobatidae) from northern Australia and New Guinea. Ichthyological Research, 63(1), 94–109. 10.1007/s10228-015-0480-9

White, W. T., O’Neill, H. L., & Naylor, G. J. P. (2022). Chapter 2: Taxonomy and Diversity of Extant Elasmobranchs. In Biology of Sharks and Their Relatives (3rd edn). CRC Press.

Winton, M., Natanson, L., Kneebone, J., Cailliet, G., & Ebert, D. (2013). Life history of Bathyraja trachura from the eastern Bering Sea, with evidence of latitudinal variation in a deep-sea skate species. Journal of the Marine Biological Association of the United Kingdom, 94, 411–422. 10.1017/S0025315413001525

