## Supplementary Material for "The allometry of spiracles and gill slits: a window into the respiratory capacity of rays"

Contents

Table S1. Species included in the analysis with the number of individuals (N).

Table S2. Linear model formulas to evaluate isometry for each swimming style.

Table S3. Species whose taxonomic names changed since the publication of the phylogenetic trees and *Rays of the World*.

Table S4. Species included in the phylogenetic trees that were excluded from analysis and the reason for exclusion.

Fig. S1. Model coefficient estimates of spiracle length and spiracle surface area across 88 species.

Fig. S2. Spiracle length and spiracle surface area across 88 species.

Fig. S3. Relative spiracle length and total length across 66 species.

Fig. S4. Relative mean gill slit height and total length across 66 species.

Fig. S5. Relative spiracle length and disc width across 22 species.

Fig. S6. Relative relative mean gill slit height and disc width across 22 species.

Table S1. Species included in the analysis with the number of individuals (N).

| Order | Family | Species | N |
|---|---|---|---|
| Myliobatiformes | Dasyatidae | *Bathytoshia lata* | 7 |
| Myliobatiformes | Dasyatidae | *Dasyatis chrysonota* | 8 |
| Myliobatiformes | Dasyatidae | *Dasyatis pastinaca* | 4 |
| Myliobatiformes | Dasyatidae | *Fontitrygon margarita* | 5 |
| Myliobatiformes | Dasyatidae | *Himantura leoparda* | 4 |
| Myliobatiformes | Dasyatidae | *Himantura uarnak* | 4 |
| Myliobatiformes | Dasyatidae | *Maculabatis gerrardi* | 5 |
| Myliobatiformes | Dasyatidae | *Neotrygon caeruleopunctata* | 5 |
| Myliobatiformes | Dasyatidae | *Pastinachus sephen* | 4 |
| Myliobatiformes | Dasyatidae | *Pteroplatytrygon violacea* | 4 |
| Myliobatiformes | Dasyatidae | *Taeniura lymma* | 7 |
| Myliobatiformes | Dasyatidae | *Telatrygon zugei* | 5 |
| Myliobatiformes | Gymnuridae | *Gymnura natalensis* | 5 |
| Myliobatiformes | Gymnuridae | *Gymnura poecilura* | 13 |
| Myliobatiformes | Gymnuridae | *Gymnura tentaculata* | 5 |
| Myliobatiformes | Gymnuridae | *Gymnura zonura* | 5 |
| Myliobatiformes | Myliobatidae | *Aetomylaeus bovinus* | 9 |
| Myliobatiformes | Myliobatidae | *Aetomylaeus wafickii* | 7 |
| Myliobatiformes | Myliobatidae | *Myliobatis aquila* | 10 |
| Myliobatiformes | Plesiobatidae | *Plesiobatis daviesi* | 4 |
| Myliobatiformes | Potamotrygonidae | *Potamotrygon orbignyi* | 5 |
| Myliobatiformes | Rhinopteridae | *Rhinoptera marginata* | 5 |
| Myliobatiformes | Urolophidae | *Urolophus deforgesi* | 5 |
| Myliobatiformes | Urolophidae | *Urolophus neocaledoniensis* | 5 |
| Myliobatiformes | Urolophidae | *Urolophus papilio* | 5 |
| Myliobatiformes | Urotrygonidae | *Urobatis halleri* | 5 |
| Myliobatiformes | Urotrygonidae | *Urobatis jamaicensis* | 4 |
| Myliobatiformes | Urotrygonidae | *Urotrygon chilensis* | 7 |
| Myliobatiformes | Urotrygonidae | *Urotrygon munda* | 5 |
| Myliobatiformes | Urotrygonidae | *Urotrygon rogersi* | 5 |
| Rajiformes | Anacanthobatidae | *Anacanthobatis marmorata* | 8 |
| Rajiformes | Arhynchobatidae | *Arctoraja parmifera* | 12 |
| Rajiformes | Arhynchobatidae | *Atlantoraja castelnaui* | 5 |
| Rajiformes | Arhynchobatidae | *Bathyraja aleutica* | 5 |

| | | | |
|---|---|---|---|
| Rajiformes | Arhynchobatidae | *Bathyraja interrupta* | 10 |
| Rajiformes | Arhynchobatidae | *Bathyraja kincaidii* | 4 |
| Rajiformes | Arhynchobatidae | *Bathyraja multispinis* | 4 |
| Rajiformes | Arhynchobatidae | *Bathyraja richardsoni* | 6 |
| Rajiformes | Arhynchobatidae | *Bathyraja spinicauda* | 5 |
| Rajiformes | Arhynchobatidae | *Psammobatis normani* | 4 |
| Rajiformes | Arhynchobatidae | *Psammobatis rudis* | 5 |
| Rajiformes | Arhynchobatidae | *Psammobatis scobina* | 5 |
| Rajiformes | Arhynchobatidae | *Sympterygia bonapartii* | 5 |
| Rajiformes | Gurgesiellidae | *Cruriraja hulleyi* | 10 |
| Rajiformes | Rajidae | *Amblyraja georgiana* | 6 |
| Rajiformes | Rajidae | *Amblyraja radiata* | 7 |
| Rajiformes | Rajidae | *Beringraja binoculata* | 6 |
| Rajiformes | Rajidae | *Caliraja rhina* | 8 |
| Rajiformes | Rajidae | *Dipturus nidarosiensis* | 4 |
| Rajiformes | Rajidae | *Leucoraja erinacea* | 4 |
| Rajiformes | Rajidae | *Leucoraja wallacei* | 10 |
| Rajiformes | Rajidae | *Okamejei kenojei* | 4 |
| Rajiformes | Rajidae | *Raja brachyura* | 4 |
| Rajiformes | Rajidae | *Raja ocellifera* | 10 |
| Rajiformes | Rajidae | *Raja straeleni* | 8 |
| Rajiformes | Rajidae | *Rajella leoparda* | 11 |
| Rajiformes | Rajidae | *Rostroraja alba* | 5 |
| Rhinopristiformes | Glaucostegidae | *Glaucostegus cemiculus* | 7 |
| Rhinopristiformes | Glaucostegidae | *Glaucostegus thouin* | 8 |
| Rhinopristiformes | Pristidae | *Anoxypristis cuspidata* | 5 |
| Rhinopristiformes | Rhinidae | *Rhina ancylostomus* | 12 |
| Rhinopristiformes | Rhinobatidae | *Acroteriobatus andysabini* | 6 |
| Rhinopristiformes | Rhinobatidae | *Acroteriobatus annulatus* | 11 |
| Rhinopristiformes | Rhinobatidae | *Acroteriobatus leucospilus* | 8 |
| Rhinopristiformes | Rhinobatidae | *Pseudobatos glaucostigma* | 4 |
| Rhinopristiformes | Rhinobatidae | *Pseudobatos leucorhynchus* | 8 |
| Rhinopristiformes | Rhinobatidae | *Pseudobatos percellens* | 5 |
| Rhinopristiformes | Rhinobatidae | *Rhinobatos albomaculatus* | 8 |
| Rhinopristiformes | Rhinobatidae | *Rhinobatos irvinei* | 7 |
| Rhinopristiformes | Rhinobatidae | *Rhinobatos rhinobatos* | 7 |
| Rhinopristiformes | Rhinobatidae | *Rhinobatos schlegelii* | 4 |

| | | | |
|---|---|---|---|
| Rhinopristiformes | Zanobatidae | *Zanobatus schoenleinii* | 4 |
| Torpediniformes | Narcinidae | *Benthobatis marcida* | 5 |
| Torpediniformes | Narcinidae | *Benthobatis moresbyi* | 5 |
| Torpediniformes | Narcinidae | *Discopyge tschudii* | 6 |
| Torpediniformes | Narcinidae | *Narcine bancroftii* | 5 |
| Torpediniformes | Narcinidae | *Narcine brasiliensis* | 8 |
| Torpediniformes | Narcinidae | *Narcine entemedor* | 4 |
| Torpediniformes | Narcinidae | *Narcine prodorsalis* | 4 |
| Torpediniformes | Narcinidae | *Narcine rierai* | 5 |
| Torpediniformes | Narkidae | *Narke dipterygia* | 5 |
| Torpediniformes | Narkidae | *Temera hardwickii* | 4 |
| Torpediniformes | Torpedinidae | *Tetronarce californica* | 9 |
| Torpediniformes | Torpedinidae | *Tetronarce cowleyi* | 8 |
| Torpediniformes | Torpedinidae | *Tetronarce nobiliana* | 4 |
| Torpediniformes | Torpedinidae | *Torpedo fuscomaculata* | 13 |
| Torpediniformes | Torpedinidae | *Torpedo sinuspersici* | 8 |
| Torpediniformes | Torpedinidae | *Torpedo torpedo* | 7 |

Table S2. Linear model formulas to evaluate isometry for each swimming style.

| **Swimming Style** | **Model formula** |
|---|---|
| Caudal Propulsion | $log_{10}$(relative gill slit height) ~ $log_{10}$(total length) + (1 + log10(disc width) \| gr(species)) |
| | $log_{10}$(relative spiracle length) ~ $log_{10}$(total length) + (1 + log10(disc width) \| gr(species)) |
| Pectoral Oscillation | $log_{10}$(relative gill slit height) ~ $log_{10}$(disc width) + (1 + log10(disc width) \| gr(species)) |
| | $log_{10}$(relative spiracle length) ~ $log_{10}$(disc width) + (1 + log10(disc width) \| gr(species)) |
| Pectoral Undulation | $log_{10}$(relative gill slit height) ~ $log_{10}$(disc width) + (1 + log10(disc width) \| gr(species)) |
| | $log_{10}$(relative spiracle length) ~ $log_{10}$(disc width) + (1 + log10(disc width) \| gr(species)) |

Table S3. Species whose taxonomic names changed since the publication of the phylogenetic trees (Stein et al. 2018) and *Rays of the World* (Last et al. 2016).

| **Old Species Name** | **Description of Change** | **New Species Name** |
|---|---|---|
| *Rhynchobatus compagnoi* | Synonym | *Rhynchobatus cooki* |
| *Myliobatis californicus* | Spelling updated | *Myliobatis californica* |
| *Hemitrygon bennetti* | Spelling updated | *Hemitrygon bennettii* |
| *Myliobatis peruvianus* | Spelling updated | *Myliobatis peruviana* |
| *Myliobatis tenuicaudatus* | Spelling updated | *Myliobatis tenuicaudata* |
| *Potamotrygon schuhmacheri* | Spelling updated | *Potamotrygon schuhemacheri* |
| *Amblyraja geogiana* | Spelling updated | *Amblyraja georgiana* |
| *Bathyraja parmifera* | Genus updated | *Arctoraja parmifera* |
| *Bathyraja pathera* | Spelling updated | *Bathyraja panthera* |
| *Bathyraja simoterus* | Genus updated | *Arctoraja simoterus* |
| *Beringraja cortezensis* | Genus updated | *Caliraja cortezensis* |
| *Beringraja inornata* | Genus updated | *Caliraja inornata* |
| *Beringraja rhina* | Genus updated | *Caliraja rhina* |
| *Beringraja stellulata* | Genus updated | *Caliraja stellulata* |
| *Dentiraja oculata* | Spelling updated | *Dentiraja oculus* |
| *Dipturus macrocaudus* | Spelling updated | *Dipturus macrocauda* |
| *Rioraja agassizi* | Spelling updated | *Rioraja agassizii* |
| *Schroederobatis* | Species name missing | *Schroederobatis americana* |
| *Zearaja argentinensis* | Genus updated | *Dipturus argentinensis* |
| *Torpedo suessi* | Spelling updated | *Torpedo suessii* |

Table S4. Species included in the phylogenetic trees (Stein et al. 2018) that were excluded from analysis and the reason for exclusion.

| **Species** | **Reason for exclusion** |
|---|---|
| *Rhinoraja longicauda* | Synonym of *Bathyraja longicauda* |
| *Anacanthobatis donghaiensis* | Synonym of *Sinobatis borneensis* |
| *Anacanthobatis nanhaiensis* | Synonym of *Sinobatis borneensis* |
| *Dipturus flavirostris* | Synonym of *Zearaja chilensis* |
| *Okamejei porosa* | Synonym of *Okamejei kenojei* |
| *Raja atriventralis* | Synonym of *Okamejei kenojei* |
| *Raja macrorynchus* | Synonym of *Dipturus batis* |
| *Raja rondeleti* | Synonym of *Leucoraja fullonica* |
| *Raja rouxi* | Synonym of *Raja straeleni* |
| *Leucoraja caribbaea* | Synonym of *Leucoraja garmani* |
| *Amblyraja robertsi* | Synonym of *Amblyraja hyperborea* |
| *Amblyraja badia* | Synonym of *Amblyraja hyperborea* |
| *Rajella alia* | Synonym of *Rostroraja ackleyi* |
| *Leucoraja virginica* | Synonym of *Leucoraja garmani* |
| *Narcine bicolor* | Questionably valid |
| *Narcine nigra* | Questionably valid |
| *Torpedo microdiscus* | Synonym of *Tetronarce tremens* |
| *Torpedo fairchildi* | Synonym of *Tetronarce nobiliana* |
| *Torpedo semipelagica* | Synonym of *Tetronarce tremens* |
| *Torpedo zugmayeri* | Synonym of *Torpedo sinuspersici* |
| *Torpedo peruana* | Synonym of *Tetronarce tremens* |
| *Torpedo macneilli* | Synonym of *Tetronarce nobiliana* |
| *Typhlonarke tarakea* | Synonym of *Typhlonarke aysoni* |
| *Heteronarce prabhui* | Synonym of *Hetronarce mollis* |
| *Crassinarke dormitor* | Synonym of *Narke japonica* |
| *Trygonorrhina melaleuca* | Synonym of *Trygonorrhina dumerilii* |
| *Rhinobatos formosensis* | Synonym of *Rhinobatos schlegelii* |
| *Gymnura crooki* | Synonym of *Gymnura poecilura* |
| *Gymnura bimaculata* | Synonym of *Gymnura japonica* |
| *Urotrygon serrula* | Synonym of *Utrotygon munda* |
| *Potamotrygon dumerilii* | Synonym of *Potamotrygon orbignyi* |
| *Potamotrygon castexi* | Synonym of *Potamotrygon falkneri* |
| *Dasyatis gigantea* | Synonym of *Bathytoshia brevicaudata* |

| | |
|---|---|
| *Dasyatis matsubarai* | Synonym of *Bathytoshia brevicaudata* |
| *Dasyatis multispinosa* | Synonym of *Bathytoshia brevicaudata* |
| *Dasyatis thetidis* | Synonym of *Bathytoshia lata* |
| *Dasyatis ushiei* | Synonym of *Bathytoshia lata* |
| *Myliobatis rhombus* | Questionably valid |
| *Myliobatis australis* | Synonym of *Myliobatis tenuicaudata* |

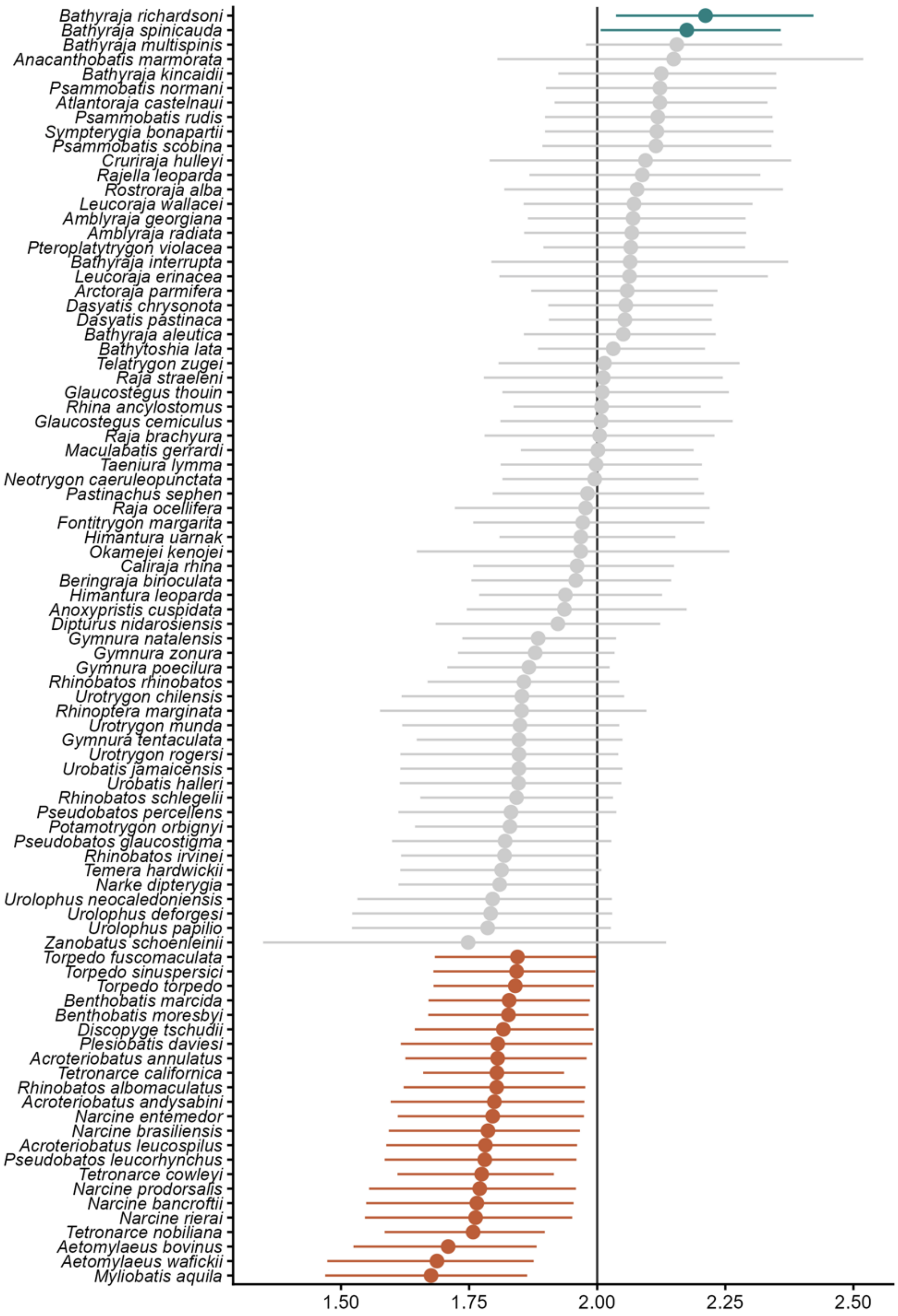


Fig. S1. Model coefficient estimates of spiracle length and spiracle surface area across 88 species. Lines for each coefficient indicate the 95% credible interval. Color indicates the statistical significance based on the confidence interval: above 2 (turquoise), not different from 2 (grey), and below 2 (rust).

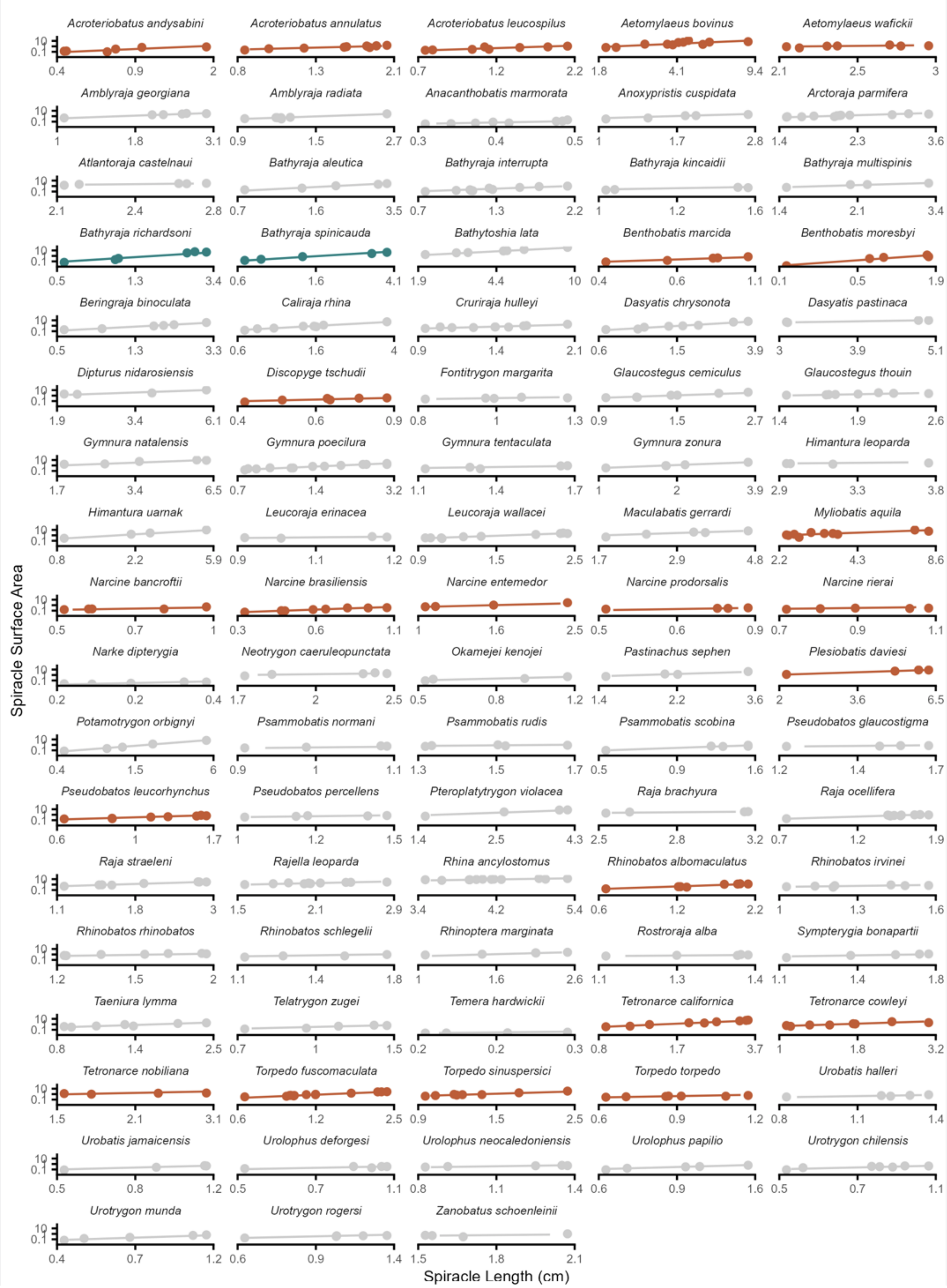


Fig. S2. Spiracle length and spiracle surface area across 88 species. Lines are the mean posterior draw for each species from the Bayesian mixed model. Color indicates the statistical significance of the coefficient (slope estimate) based on the confidence interval (see Fig. S2): above 2 (turquoise), not different from 2 (grey), and below 2 (rust).

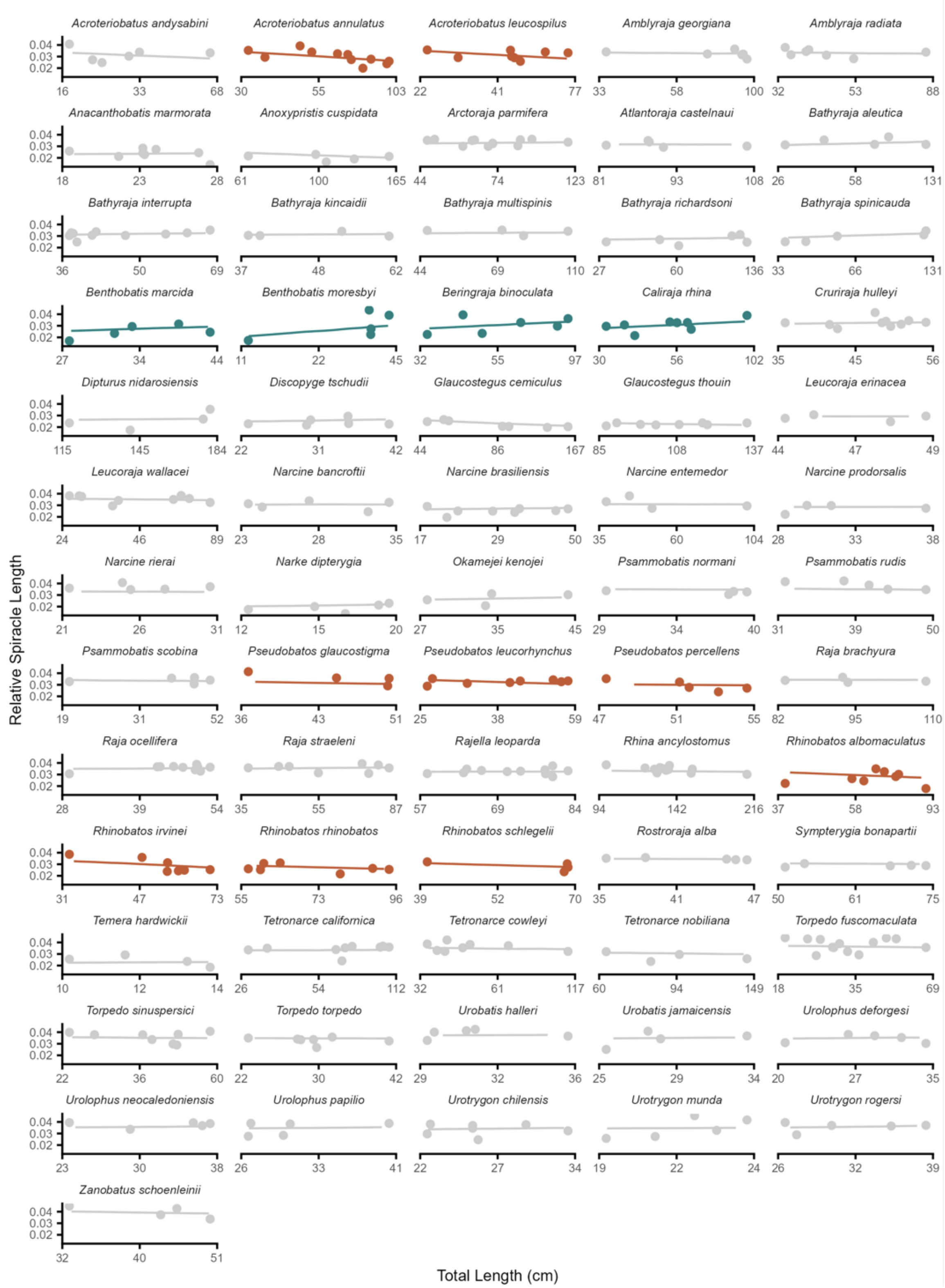


Fig. S3. Relative spiracle length and total length across 66 species. Lines are the mean posterior draw for each species from the Bayesian mixed model. Color indicates the statistical significance of the coefficient (slope estimate) based on the confidence interval (see Fig. S4): above 0 (turquoise), not different from 0 (grey), and below 0 (rust).

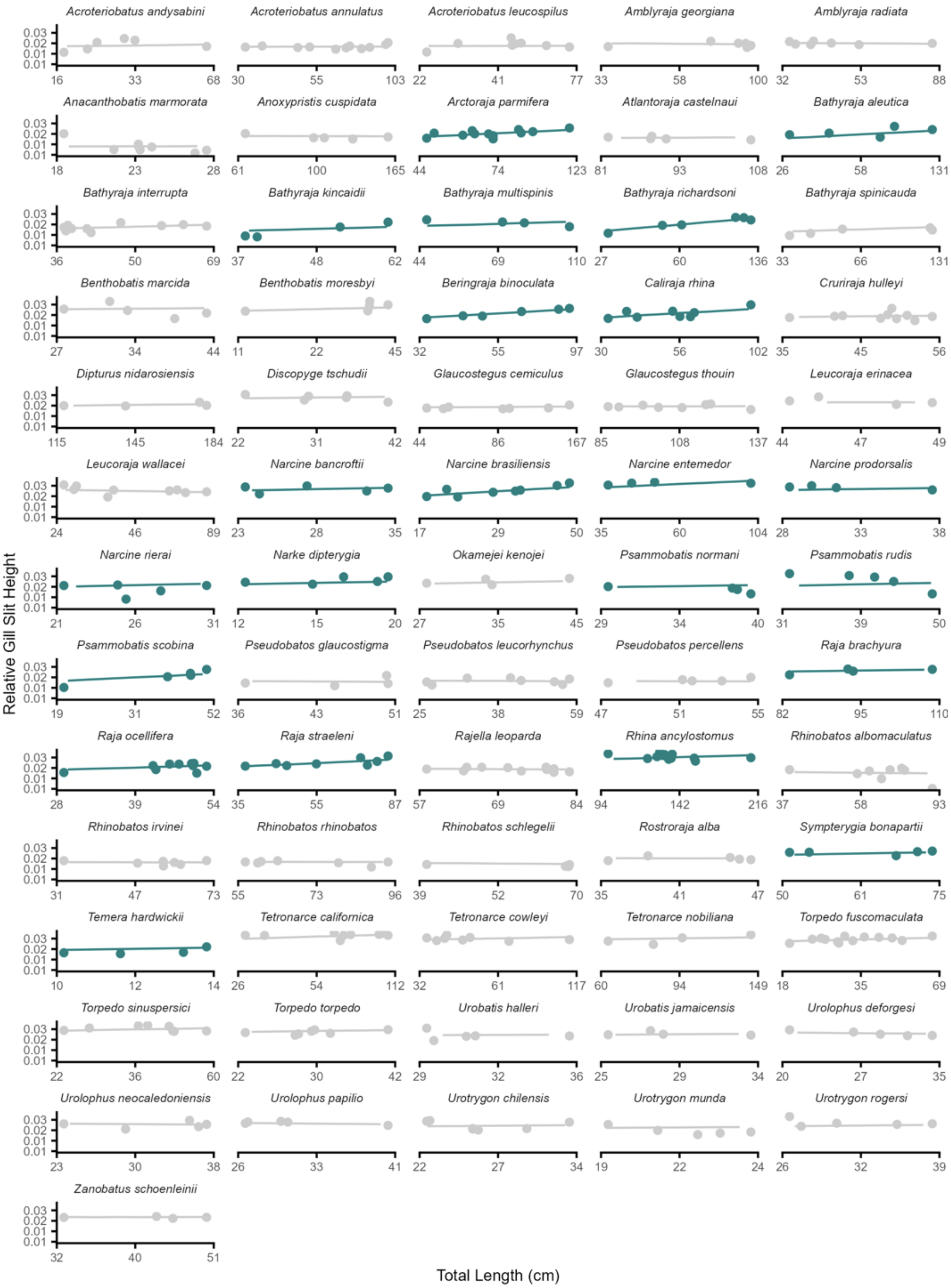


Fig. S4. Relative mean gill slit height and total length across 66 species. Lines are the mean posterior draw for each species from the Bayesian mixed model. Color indicates the statistical significance of the coefficient (slope estimate) based on the confidence interval (see Fig. S6): above 0 (turquoise) and not different from 0 (grey).

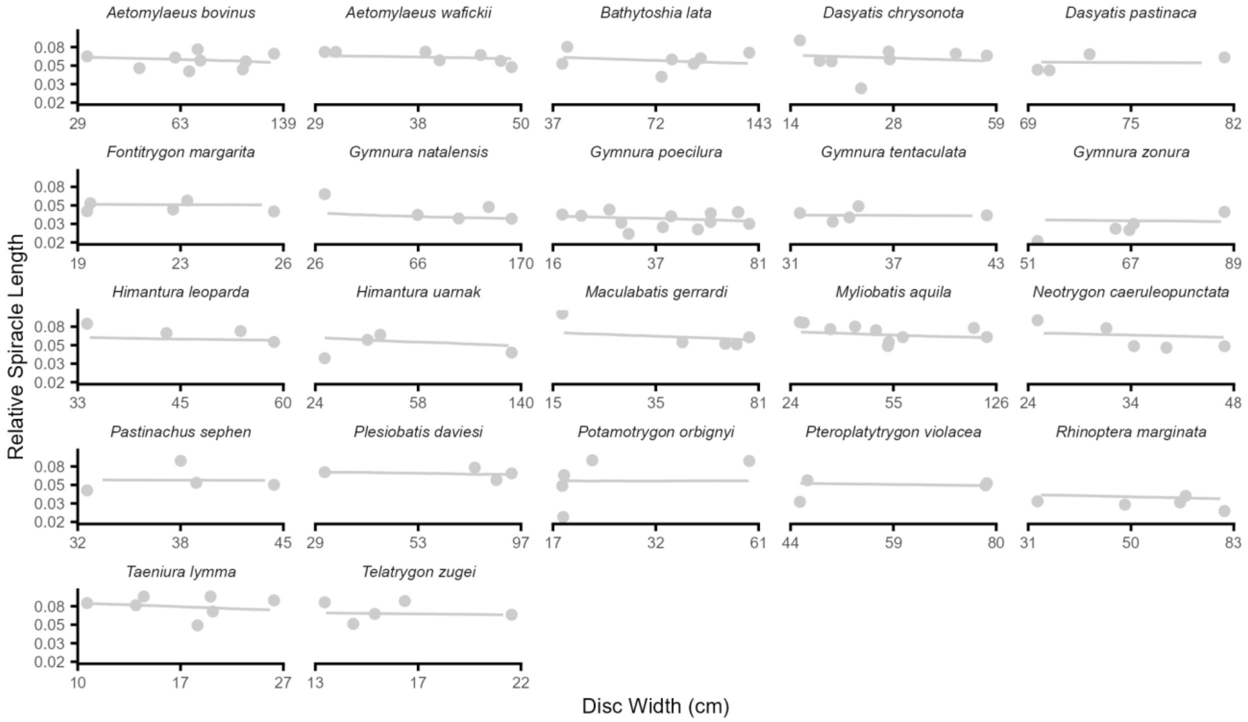


Fig. S5. Relative spiracle length and disc width across 22 species. Lines are the mean posterior draw for each species from the Bayesian mixed model. Grey color indicates the statistical significance of the coefficient (slope estimate) based on the confidence interval not being different from 0 (see Fig. S8).

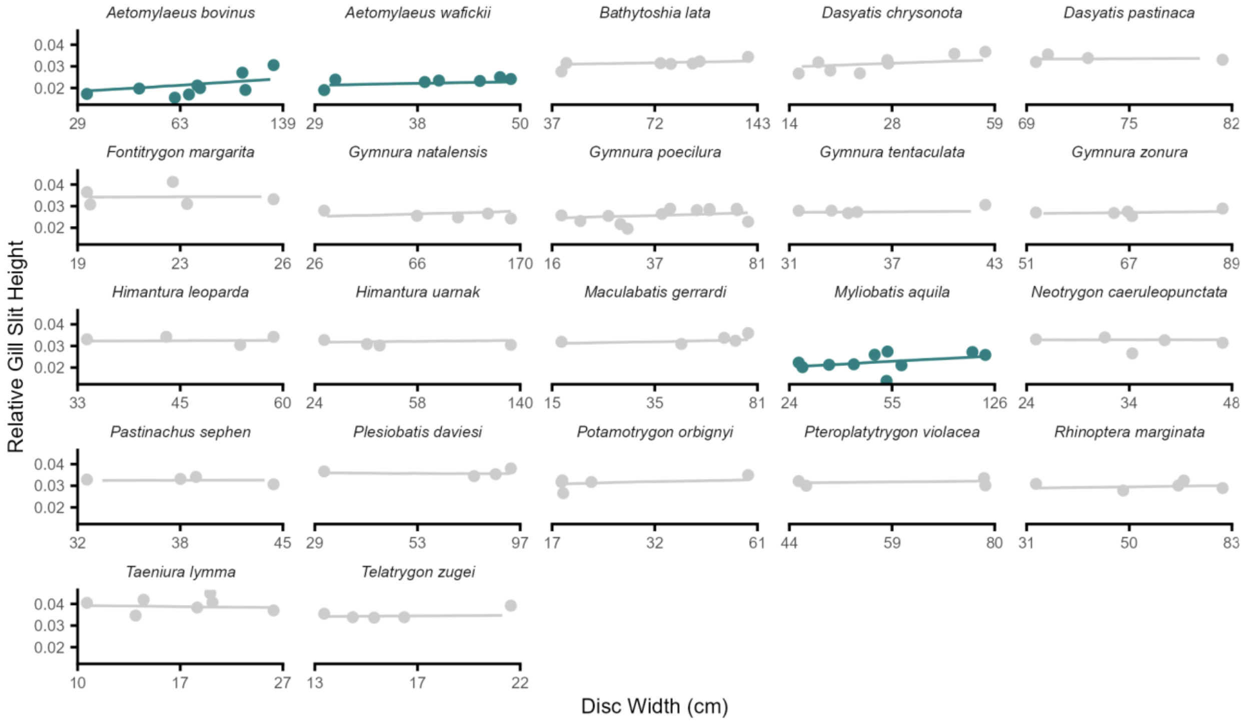


Fig. S6. Relative relative mean gill slit height and disc width across 22 species. Lines are the mean posterior draw for each species from the Bayesian mixed model. Color indicates the statistical significance of the coefficient (slope estimate) based on the confidence interval (see Fig. S10): above 0 (turquoise) and not different from 0 (grey).
